# Translational control of lipogenesis links protein synthesis and phosphoinositide signaling with nuclear division

**DOI:** 10.1101/2021.01.03.425130

**Authors:** Nairita Maitra, Staci Hammer, Clara Kjerfve, Vytas A. Bankaitis, Michael Polymenis

## Abstract

Continuously dividing cells coordinate their growth and division. How fast cells grow in mass determines how fast they will multiply. Yet, there are few, if any, examples of a metabolic pathway that actively drives a cell cycle event instead of just being required for it. Here, we show that translational upregulation of lipogenic enzymes in yeast increased the abundance of lipids and accelerated nuclear elongation and division. De-repressing translation of acetyl CoA carboxylase and fatty acid synthase also suppressed cell cycle-related phenotypes, including delayed nuclear division, associated with Sec14p phosphatidylinositol transfer protein deficiencies, and the irregular nuclear morphologies of mutants defective in phosphatidylinositol 4-OH kinase activities. Our results show that increased lipogenesis drives a critical cell cycle landmark and report a phosphoinositide signaling axis in control of nuclear division. The broad conservation of these lipid metabolic and signaling pathways raises the possibility these activities similarly govern nuclear division in mammals.

## INTRODUCTION

The sequential and essential action of acetyl-CoA carboxylase (Acc1p in yeast) and fatty acid synthase (FAS) make fatty acids in all organisms. Acc1p carboxylates acetyl-coenzyme A (CoA) to make malonyl-CoA (Al-Feel et al., 1992), which is subsequently consumed by FAS as the building block for fatty acid synthesis. Yeast FAS is made of two different polypeptides, α-(Fas2p) and β-subunits (Fas1p), in a 2.6 MDa α_6_β_6_ dodecamer (Lomakin et al., 2007; Leibundgut et al., 2008; Jenni et al., 2007). The correct Fas1p and Fas2p stoichiometry is achieved by a mechanism where Fas1p controls *FAS2* mRNA levels (Wenz, 2001). Moreover, Fas1p initiates FAS assembly via a co-translational interaction with nascent Fas2p (Shiber et al., 2018; Fischer et al., 2020). The multiple catalytic activities of FAS iteratively add two carbons at a time from a malonyl-CoA donor to an acetyl-CoA acceptor. The reaction terminates after seven or eight cycles, generating 16-or 18-carbon fatty acids, respectively (Lomakin et al., 2007; Jenni et al., 2007).

There is intense contemporary interest in lipogenesis during cell division given the implications for cell proliferation (Storck et al., 2018). Higher levels of lipogenic enzymes are a near-universal marker of tumors (Kuhajda et al., 1994; Baenke et al., 2013; Beloribi-Djefaflia et al., 2016), and FAS is one of the few metabolic enzymes targeted in cancer clinical trials (NCT02595372, NCT02980029). Three lines of evidence suggest that *de novo* lipid synthesis impinges on events late in the cell cycle. First, the translational efficiency of acetyl-CoA carboxylase and fatty acid synthase mRNA transcripts peaks in mitosis (Blank et al., 2017b; a). Second, the abundance of lipids similarly peaks late in the cell cycle (Blank et al., 2020). The midbodies in human cells, where cleavage occurs at the end of the cell cycle, have a distinct lipid composition (Atilla-Gokcumen et al., 2014). Third, *de novo* lipid synthesis is essential for mitosis in humans (Scaglia et al., 2014) and yeast (Al-Feel et al., 2003; Schneiter et al., 1996).

Nuclear membrane dynamics are pronounced during mitosis. For example, there is a dramatic proliferation in new nuclear membranes during mitosis. This proliferation occurs irrespective of whether the mitosis is open (i.e., the nuclear envelope disassembles in mitosis as in animal cells) or whether it is closed (i.e., nuclear envelope integrity is maintained throughout mitosis as in fungi). In fission and budding yeast, loss-of-function mutations in lipid synthesis perturb the size and shape of the nuclear envelope (Witkin et al., 2012; Walters et al., 2012, 2014; Siniossoglou, 2013; Santos-Rosa et al., 2005; Zach and Prevorovsky, 2018; Kume et al., 2017). The evidence that lipid synthesis is required for cell division notwithstanding, it remains unclear whether lipid biogenesis promotes specific cell cycle events and, if so, what those events might be.

Sec14p is the founding member of a major family of phosphatidylinositol (PtdIns) transfer proteins (PITPs). Sec14p and other PITPs stimulate the activities of PtdIns 4-OH kinases by rendering PtdIns a superior substrate for these enzymes – thereby potentiating PtdIns(4)P-dependent signaling (Wang et al., 2019). Sec14p executes an essential cellular activity in promoting membrane trafficking from yeast Golgi/endosomal compartments, but this cellular requirement is obviated when lipid metabolism is appropriately altered. For example, inactivation of CDP-choline salvage pathway for PtdCho synthesis relieves cells of the essential Sec14p requirement (Cleves et al., 1991). In addition to membrane trafficking, Sec14p also controls cell cycle progression (Huang et al., 2018). Reductions in Sec14p activity of insufficient magnitude to compromise cell viability nonetheless result in aberrantly larger cells that progress through the cell cycle more slowly with notable delays in progression through the G2/M phases (Huang et al., 2018). How Sec14p impinges on cell cycle progression and how it is functionally integrated with lipid synthesis in the cell cycle remains poorly understood.

Herein, we demonstrate that coordinate de-repression of *ACC1* and *FAS1* translation results in elevated lipogenesis that promotes nuclear division in yeast. Furthermore, we report that loss-of-function mutations in the conserved Sec14p delay nuclear division, and that de-repressing translational control of *ACC1* and *FAS1* not only corrects the delayed nuclear division associated with reduced Sec14 activity, but suppresses other mitotic phenotypes associated with Sec14p insufficiencies. Likewise, we document that loss of phosphatidylinositol 4-OH kinase activity deranges nuclear morphology, and that these phenotypes too are rescued by enhanced translation of *ACC1* and *FAS1* mRNAs. Taken together, these results identify translational control as a critical input for tuning lipid metabolism to progression through the cell cycle. The data also indicate that lipid synthesis is not only required for mitosis, but that it actively promotes nuclear division -- a key landmark of the eukaryotic cell cycle.

## RESULTS

### Translational control elements in lipogenic enzymes

The translational efficiency of acetyl-CoA carboxylase and fatty acid synthase is upregulated late in the cell cycle and, in that regard, removing a uORF from the 5’-leader of the *ACC1* transcript de-represses the translation of *ACC1* (Blank et al., 2017b). The 5’-leader of *FAS1* is 551 nt-long (Xu et al., 2009), and has two uORFs (Figure 1A). Ribosome profiling experiments report that both *FAS1* uORFs are likely translated in *S. cerevisiae* (Ingolia et al., 2009). The longer and more distal uORF initiates from a non-AUG start codon. A uORF at the same position, albeit of varying length and sequence, is present in most *Saccharomyces* genomes, except in *S. mikatae*, where the AAG start codon is mutated to AAA (Figure 1 - figure supplement 1, top). The proximal uORF is short, encoding a six amino acid peptide, and it is well-conserved in the *Saccharomyces* genus (Figure 1 - figure supplement 1, bottom). There was a Tyr→Phe change in *S. kudriavzevii*, and a Phe→Leu change in *S. bayanus* (Figure 1 - figure supplement 1, bottom).

**FIGURE 1.**
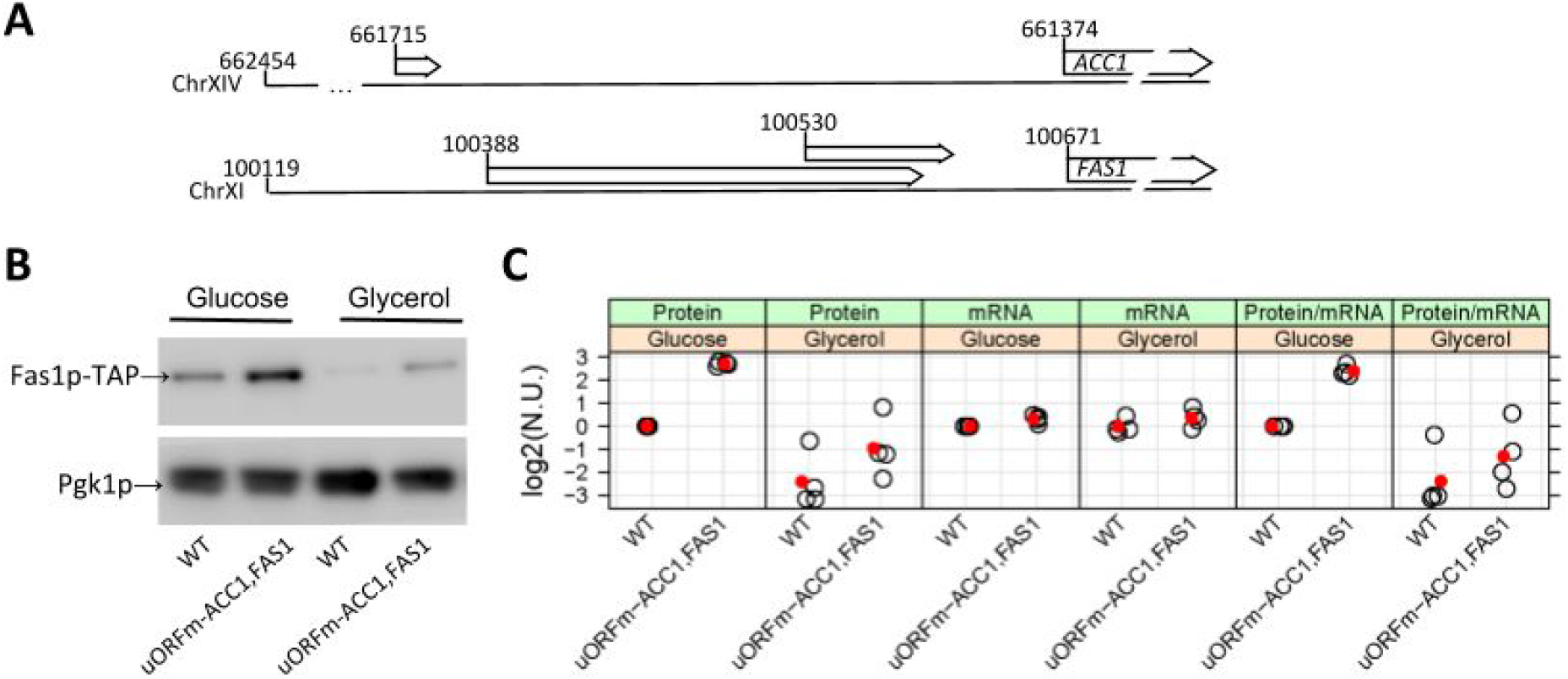
uORFs in *ACC1* and *FAS1*. A, The chromosomal coordinates of the 5’-leader sequences of the *ACC1* and *FAS1* transcripts. The 5’-ends are from (Xu et al., 2009). B, Steady-state Fas1p protein levels were measured in rich undefined media, differing in the carbon source (2% glucose or 3% glycerol) from the indicated strains carrying C-terminal TAP-tagged alleles of *FAS1* at their endogenous chromosomal locations. C, The values in the strip charts depict the relative abundance of *FAS1* mRNA and protein in *uORFm-ACC1,FAS1* cells quantified from independent experiments, from the same strains and media shown in B. Transcript levels of *FAS1* and *UBC6* were quantified by Droplet Digital PCR (ddPCR), as described in Materials and Methods. To obtain the normalized units (n.u.) on the y-axis, we first normalized for loading against the corresponding *UBC6* and Pgk1p values from the same samples. We then expressed these values as ratios against the corresponding values of wild-type *FAS1-TAP* cells, in which the uORFs are in place, from experiments run and analyzed in parallel. All the immunoblots for this figure are in file Figure 1 - source data 1, while the mRNA abundance experiments we used to generate the strip plots are in file Figure 1 - source data 2.

To test the role of translational control in fatty acid synthesis, a mutant yeast strain was generated where *cis*-elements that could repress translation of *ACC1* and *FAS1* transcripts were inactivated. Simultaneous de-repression of both *ACC1* and *FAS1* translation was expected to increase metabolic flux through *de novo* lipogenesis pathways. Because the *FAS1* uORFs are likely translated in *S. cerevisiae*, and are well conserved in other species, their start codons were mutated. The *FAS1* uORF mutations were subsequently recombined into a yeast strain carrying the *ACC1* ORF mutation we had described previously (Blank et al., 2017b). The strain carrying these three uORF mutations (*uORFm-ACC1,FAS1*) was also engineered so that the main *FAS1* ORF was TAP-tagged for purposes of protein surveillance (see Materials and Methods). As expected, asynchronous cultures of the triple uORF mutant produced elevated steady-state levels of Fas1p (Figure 1B). The ratio of the steady-state levels of the Fas1p protein over the *FAS1* mRNA was 3 to 4-fold higher in the uORF mutant cells compared to wild type cells (Figure 1C). These data were consistent with a translational de-repression mechanism for increased Fas1p expression.

To monitor Fas1p levels throughout the cell cycle, small early G1 wild-type and *uORFm-ACC1,FAS1* cells were isolated by centrifugal elutriation and Fas1p-TAP levels were examined as these cells progressed synchronously through the cell cycle (Figure 1 - figure supplement 2). Fas1p-TAP levels still oscillated in *uORFm-ACC1,FAS1* cells (Figure 1 - figure supplement 2). Whereas uORF inactivation increased the overall steady-state levels of Fas1p, the elevated abundance of Fas1p in mitotic cells was maintained. Acc1p levels were similarly periodic in the cell cycle when *ACC1* translation was de-repressed in cells lacking the *ACC1* uORF (Blank et al., 2017b). Why inactivating the uORFs does not alter the cell cycle-dependent abundance of the corresponding proteins is detailed below (see Discussion).

### De-repressing translation of *ACC1* and *FAS1* alters lipid abundances

Two approaches were used to determine whether elevated levels of Acc1p and Fas1p in *uORFm-ACC1,FAS1* cells led to increased activity of these enzymes in vivo. The first was a bioassay where the sensitivity of wild-type and triple mutant cells to cerulenin was compared. Cerulenin is a validated inhibitor of fatty acid synthase that acts by binding and inhibiting the β-ketoacyl-acyl carrier protein (ACP) synthase domain of fatty acid synthase (Omura, 1976; Price et al., 2001). Whereas proliferation of wild type cells was significantly reduced and then completely blocked at 1.5 and 2.5 μg/ml cerulenin, respectively, the *uORFm-ACC1,FAS1* cells were resistant to 2.5 μg/ml cerulenin (Figure 2 - figure supplement 1). The increased resistance of *uORFm-ACC1,FAS1* cells to cerulenin argues that translational de-repression of *ACC1* and *FAS1* leads to elevated activity of these lipogenic enzymes in vivo.

The second approach involved quantification of primary metabolites and complex lipids by GC-TOF MS, and CSH-QTOF MS/MS, respectively (See Materials and Methods). In these experiments, we examined cultures where glucose or glycerol served as sole carbon sources (Figure 2). Among the >300 metabolites that were confidently identified in cultures with glucose as sole carbon source, the alterations in metabolite abundance were almost entirely accounted for by alterations in the abundance of lipid molecular species. Increased levels of various molecular species of triglycerides (TG), phosphatidylinositol (PtdIns; PI), phosphatidylcholine (PtdCho; PC), phosphatidylethanolamine (PtdEtn; PE), and phosphatidylserine (PtdSer; PS) were recorded. A few species exhibited a lower abundance in *uORFm-ACC1,FAS1* cells and these represented the longer chain molecular species detected (PtdIns / PtdCho / PtdEtn 36:1; PtdCho 36:2; TG 50:1; TG 56:2). This pattern was reproduced in cultures where glycerol served as a carbon source although the roster of molecular species that were altered in abundance in glycerol conditions was expanded somewhat. We conclude that coordinate de-repression of *ACC1* and *FAS1* translation altered the lipidome predominantly by increasing the abundance of most lipid molecular species.

**FIGURE 2.**
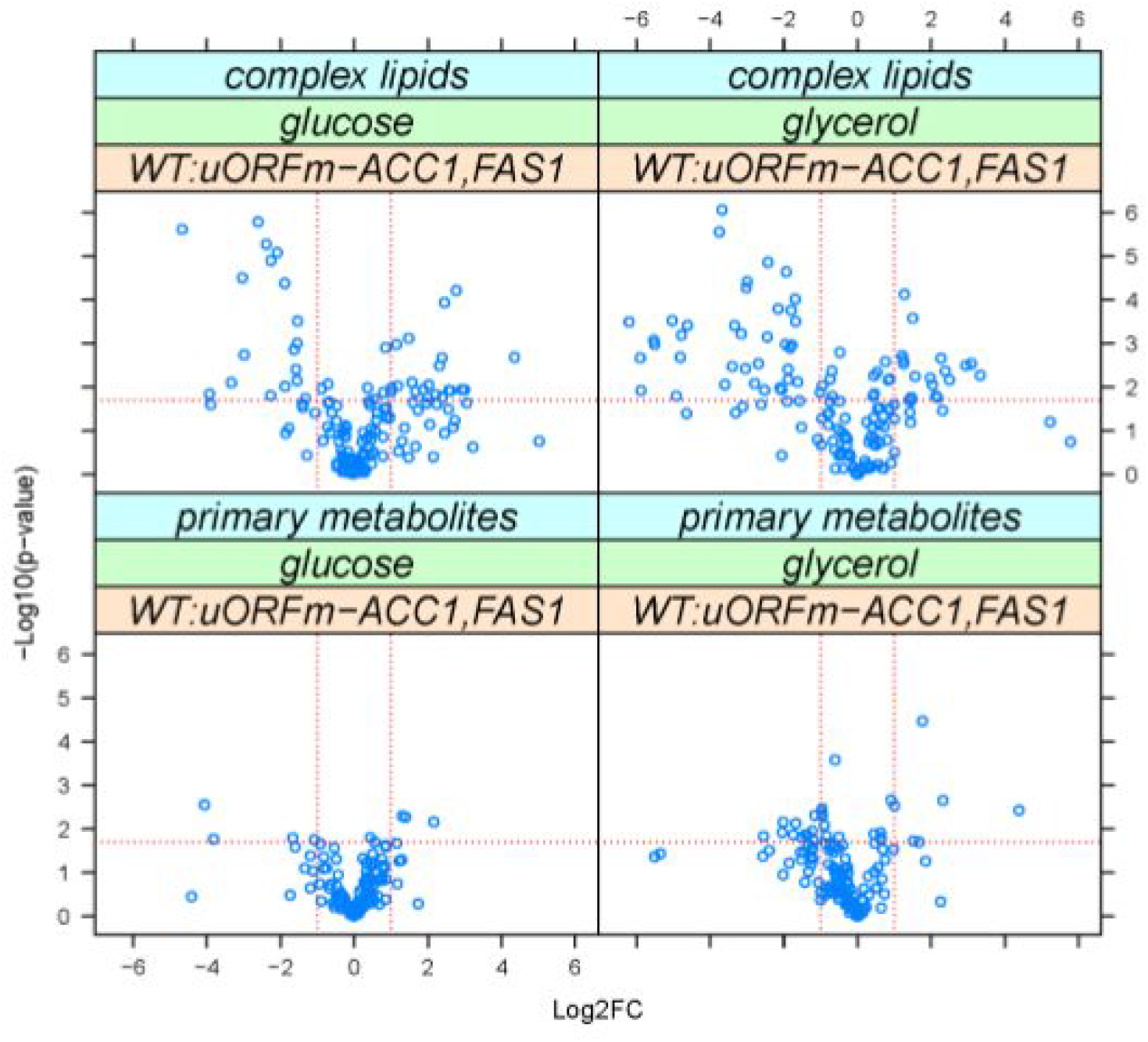
De-repressing the translational control of lipogenesis alters lipid abundances. Metabolites whose levels changed in WT vs. *uORFm-ACC1,FAS1* cells were identified from the magnitude of the difference (x-axis; Log2-fold change) and statistical significance (y-axis), indicated by the red lines. The analytical and statistical approaches are described in Materials and Methods. The values used to generate the graphs are in file Figure 2 - source data 3.

### Lipogenesis controls the timing of nuclear division

The *uORFm-ACC1,FAS1* cells represent a unique, gain-of-function context for interrogating the effects of increased lipogenesis on nuclear division. It is well-established that diminutions in lipid synthesis perturb the shape of the nuclear envelope in fungi (Witkin et al., 2012; Walters et al., 2012, 2014; Siniossoglou, 2013; Santos-Rosa et al., 2005; Zach and Prevorovsky, 2018; Kume et al., 2017). Those findings motivated examination of nuclear morphology in *uORFm-ACC1,FAS1* cells progressing synchronously through the cell cycle. As in our past studies (Blank et al., 2020, 2017b; Huang et al., 2018), centrifugal elutriation was again employed to recover highly enriched populations of un-arrested early G1 cells where the coupling between cell growth and division is maintained (Aramayo and Polymenis, 2017; Creanor and Mitchison, 1979). The timing of nuclear division was then compared for synchronous wild-type and *uORFm-ACC1,FAS1* cultures.

To that end, the nuclear envelope was visualized by immunofluorescence using the Nsp1 p component of the central core of the nuclear pore complex as a marker (see Materials and Methods and Figure 3 - figure supplement 1A). For each cell scored at one point in the cell cycle, the longest axis across the nucleus was related to the diameter of the non-elongated, spherical nuclear space and expressed as a ratio (Figure 3 - figure supplement 1A). The cell size and percentage of budded cells from the same elutriated samples were also recorded. Since yeast bud emergence marks the initiation of cell division, this event serves as a convenient proxy for marking entry into the S phase (Pringle, 1981). This landmark was therefore used to relate initiation of cell division to cell size. A key parameter that derives from such an analysis is the critical size, and this metric is defined as the size at which half the cells are budded (Soma et al., 2014). The critical size of *uORFm-ACC1,FAS1* cells was slightly smaller than that of wild type cells (Figure 3A; 36.7±1 fL vs. 40.6±2 fL, based on four independent experiments). However, compared to wild type, *uORFm-ACC1,FAS1* mutants initiated nuclear elongation and completed nuclear division disproportionately sooner – i.e., at a cell size some 10 fL smaller relative to wild type cells (Figure 3B). However, the total cell cycle time was the same for both wild type and *uORFm-ACC1,FAS1* cells (93±2 m vs. 91±5 m, p=0.3, based on the Mann-Whitney test, from at least four independent experiments in each case). Thus, although *uORFm-ACC1,FAS1* cells complete nuclear division earlier than wild-type cells do, the mutant cells delay completion of cytokinesis and cell separation. These results reveal an active and sufficient role for lipid synthesis in promoting nuclear division.

**FIGURE 3.**
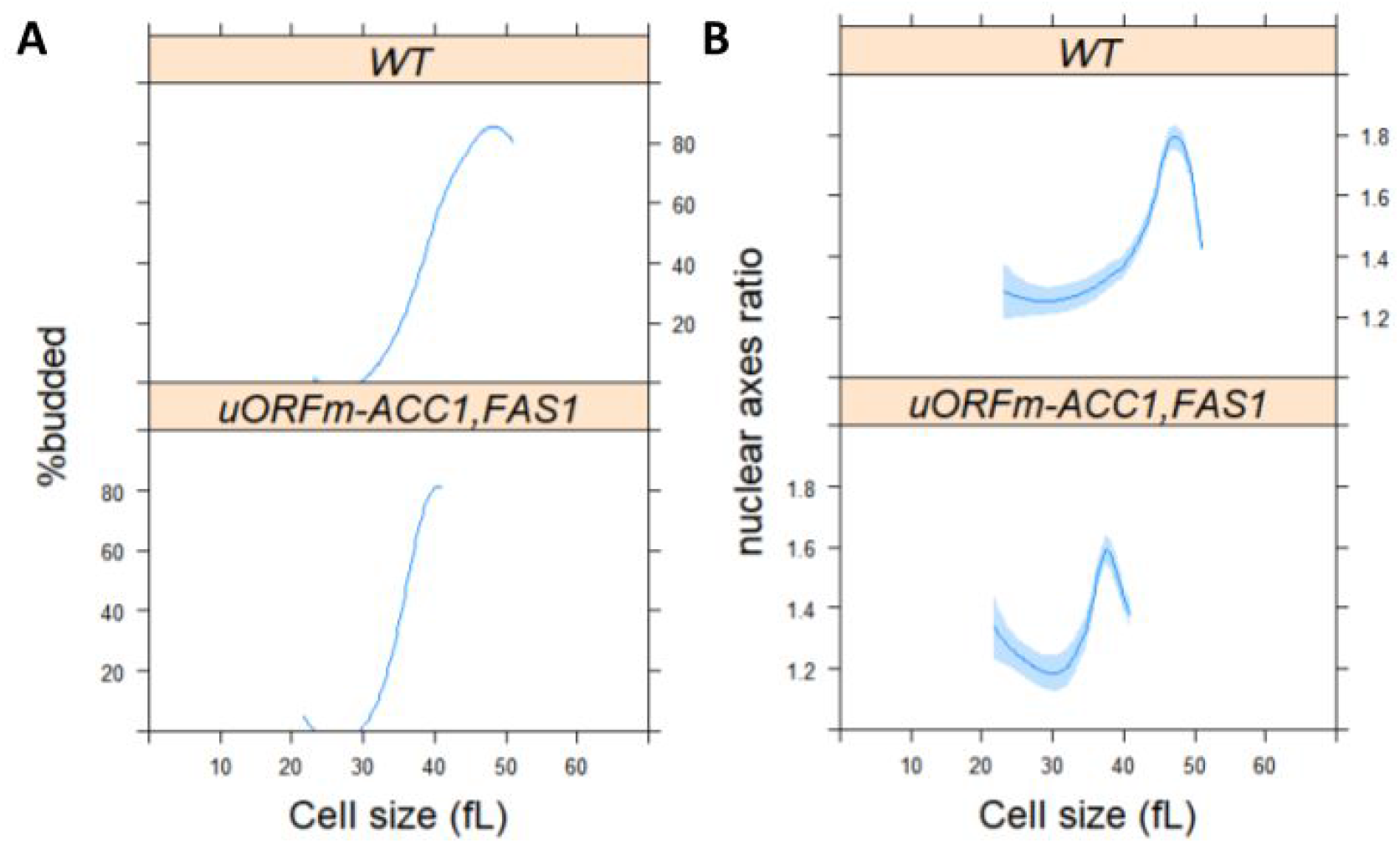
De-repressing translation of *ACC1* and *FAS1* promotes nuclear division, while inhibiting lipogenesis delays it. A, Synchronous cultures of the indicated strains and conditions were obtained by elutriation (see Materials and Methods), from which the percentage of budded cells (y-axis) is shown against the mean cell size (in fL; x-axis). B, From the same samples as in A, cells were processed for fluorescence microscopy to visualize the nucleus, as described in Materials and Methods (see also Figure 3 - figure supplement 1). Nuclear shape (y-axis) is shown against the mean cell size (in fL; x-axis), from >2,000 cells/strain. Loess curves and the std errors at a 0.95 level are shown. The values used to generate the graphs are in file Figure 3 - source data 1.

### De-repressing translation of *ACC1* and *FAS1* corrects mitotic phenotypes of *secl4* mutants

Conditional *sec14* mutants proliferate at 25°C, but not at 37°C (Novick et al., 1980). Furthermore, inhibition of lipogenesis by cerulenin exacerbates the temperature sensitivity of *sec14-1* cells (Dacquay et al., 2017). Whereas the cerulenin effect is a rather non-specific phenotype (Lee et al., 2014), this observation raised the question of whether the increased lipogenesis in *uORFm-ACC1,FAS1* cells could suppress the temperature-sensitive proliferation of *sec14-1* cells. Indeed, coordinate *ACC1* and *FAS1* uORF inactivation partially rescued growth of *sec14-1* cells at the semi-permissive temperature of 34°C (Figure 4A). This result suggests that increased lipogenesis reduced the threshold requirement of yeast cells for Sec14p.

**FIGURE 4.**
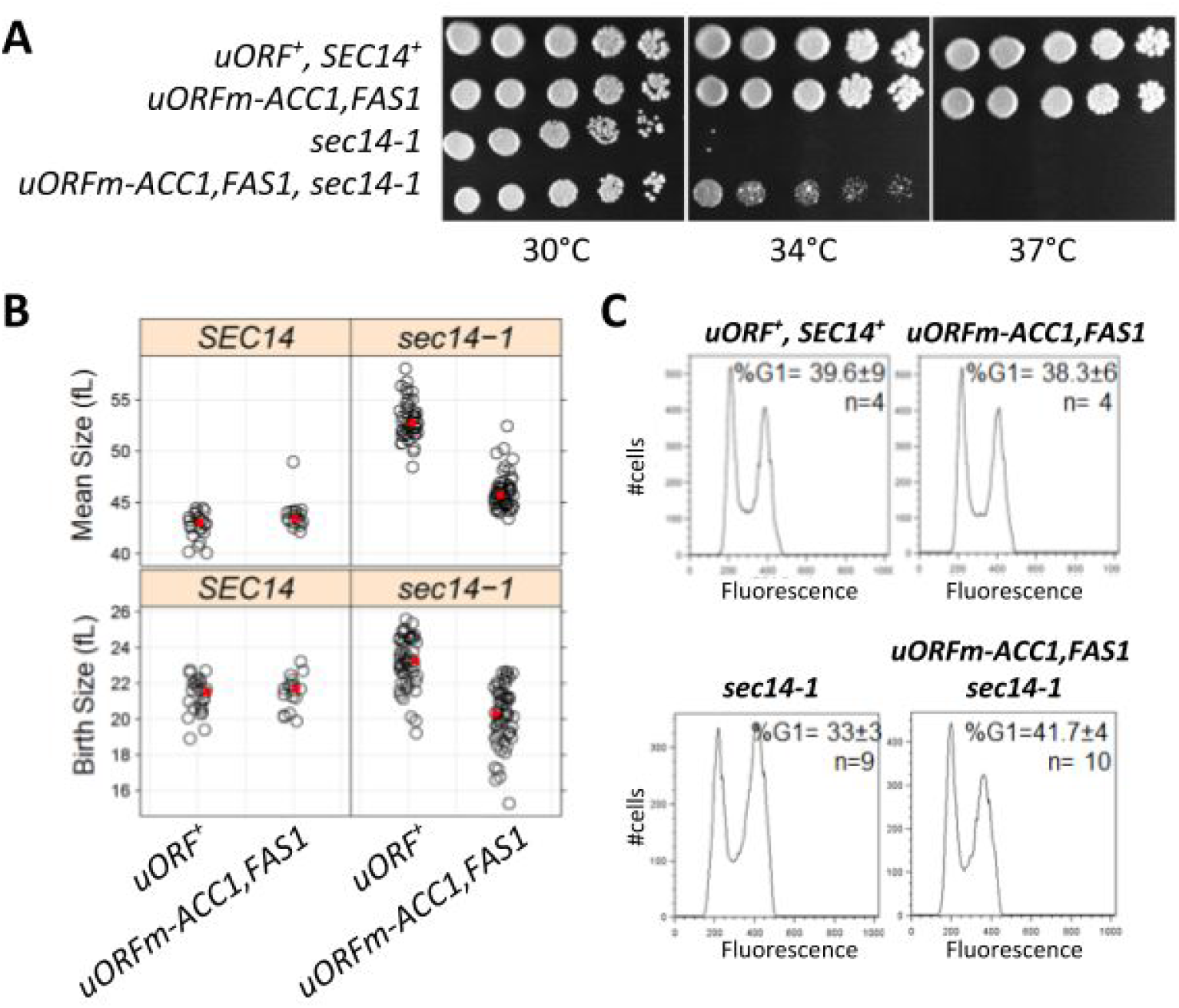
De-repressing the translational control of *ACC1* and *FAS1* suppresses the temperature sensitivity (A), large size (B), and high G2/M DNA content (C) of *sec14-1* cells. A, Serial dilutions (5-fold) of cultures of the indicated strains were spotted on solid media and grown at the temperatures shown. B, Strip plots showing the mean (top panels) and birth (bottom panels) size (y-axis) for the indicated strains. The average in each case is in red. C, Flow cytometry histograms for the indicated strains, with cell number on the y-axis and fluorescence per cell on the x-axis.

Although the Sec14p requirement for membrane trafficking through the yeast TGN/endosomal system is well-established (Bankaitis et al., 1989; Cleves et al., 1989; Nile et al., 2014), this PITP also executes cell-cycle functions. Even at the permissive temperature of 30°C *sec14-1* cells exhibit larger birth and mean sizes and are delayed in passage through the G2/M stages of the cell cycle. This latter deficiency is manifested in asynchronous cultures as increased fractions of cells in G2/M (Huang et al., 2018). Also, *sec14-1* mutants must reach a larger critical size before initiating cell division (Huang et al., 2018). Strikingly, whereas *ACC1* and *FAS1* uORF mutations did not alter the size of wild type cells, these uORF mutations completely corrected the larger birth and mean size of *sec14-1* cells (Figure 4B, right panels; p>0.05, Kruskal-Wallis, and posthoc Nemenyi tests). Moreover, the combined uORF mutations fully corrected the G2/M delay characteristic of *sec14-1ts* cells (Figure 4C). (Figure 4B, left panels). These collective results cannot be accounted for by simple models where cell enlargement is accommodated by new plasma membrane synthesis before division.

Sec14p has not been previously implicated in nuclear division. But since our data suggest that *ACC1* and *FAS1* uORF mutations correct the various cell cycle-related phenotypes of *sec14-1* cells, we examined the timing of nuclear division in synchronized cultures produced by elutriation. As we had previously reported (Huang et al., 2018), the critical size of *sec14-1* cells is larger than that of wild type cells even at the permissive temperature of 30°C (Figure 5A, compare the top two panels). Furthermore, *sec14-1* cells both initiated and completed nuclear division at a significantly larger size relative to wild-type cells (Figure 5B). Strikingly, *ACC1* and *FAS1* uORF mutations corrected both the larger critical size and the delayed nuclear division of *sec14-1* cells (Figure 5A, 5B). However, the scheduling of nuclear division in *uORFm-ACC1,FAS1, sec14-1* mutants was similar to that of wild type cells, and was not accelerated as observed in *uORFm-ACC1,FAS1* mutants (Figure 3B). These collective data not only strongly support the idea that increased lipogenesis provides for the mitotic functions of Sec14p, but also suggest that the accelerated nuclear division schedule associated with elevated lipogenesis still requires some Sec14p involvement.

**FIGURE 5.**
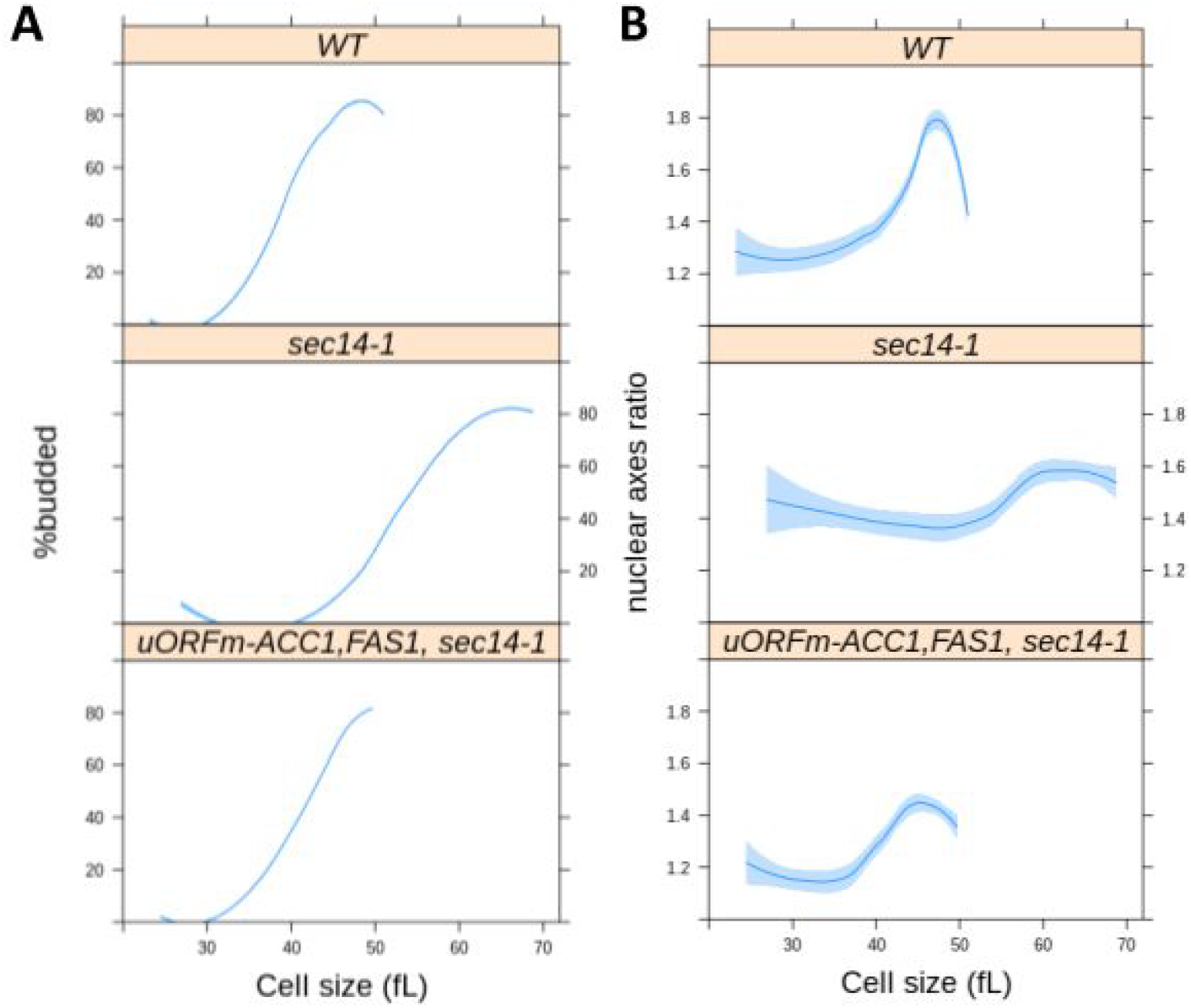
Loss-of-function *sec14-1* mutants delay nuclear division, but they are suppressed by the uORF mutations in *ACC1* and *FAS1*. From synchronous cultures of the indicated strains, the percentage of budded cells (A) and nuclear shape (B) as a function of cell size (x-axis) were measured and plotted as in Figure 3. For the wild type cells (WT), the values were the same as those in Figure 3. The values used to generate the graphs are in file Figure 5 - source data 1.

### Specificity of ‘bypass Sec14p’ mechanisms in correcting the delayed nuclear division of *sec14-1* mutants

The *ACC1* and *FAS1* uORF mutations perturb lipid metabolism (Figure 2). But fatty acids are the precursors for all glycerolipids and sphingolipids. Hence, it is difficult to reduce to a manageable level the possibilities regarding which derangements in lipid metabolism are primarily responsible for disturbed nuclear morphogenesis. Since our data point to functional interactions between the increased lipogenesis in *uORFm-ACC1,FAS1* cells and Sec14p-dependent processes (Figures 4, 5), the known roles of Sec14p were exploited as a tool that reports on distinct arms of the lipid metabolome.

As a PtdIns/PtdCho transfer protein (Grabon et al., 2019; Bankaitis et al., 1990), Sec14p leverages its heterotypic lipid binding and exchange activities to potentiate the activities of PtdIns 4-OH kinases and thereby stimulate PtdIns-4-phosphate signaling (Schaaf et al., 2008; Bankaitis et al., 2010). It is by this mechanism that Sec14p and other PITPs function as sensors of lipid metabolism and integrate the activities of distinct arms of the lipid metabolome with localized outputs of PtdIns-4-phosphate signaling (Wang et al., 2019). In that regard, Sec14p executes essential cellular activities, but the normally essential Sec14p requirement for cell viability is bypassed by derangements in specific aspects of lipid metabolism. These include loss-of-function mutations in the PtdIns-4-phosphate phosphatase Sac1 (Cleves et al., 1989; Rivas et al., 1999), the CDP-choline pathway (but not the *de novo* pathway) for PtdCho biosynthesis (Cleves et al., 1991; Xie et al., 2001), and the ergosterol and PtdIns-4-phosphate exchange protein Kes1p (Fang et al., 1996; Li et al., 2002; Mousley et al., 2012).

With regard to bypass Sec14p mutations in the CDP-choline pathway, the lethality and secretory defects of *sec14-1* mutants at 37°C are suppressed by loss of the Cki1p choline kinase which catalyzes the first reaction of this PtdCho biosynthetic pathway (Cleves et al., 1991). However, even though genetic ablation of Cki1p activity yielded cells that exhibited a slightly smaller critical size (consistent with slightly accelerated initiation of cell division), this metabolic defect failed to suppress the delayed nuclear division of *sec14-1* mutants (Figure 6). Strikingly different results were obtained for Kes1p loss-of-function mutations which we previously demonstrated correct the large cell size and cell cycle delay of *sec14-1* cells ((Huang et al., 2018); see also Figure 6A). Kes1p inactivation fully corrected the delay in the nuclear division of *sec14-1* cells (Figure 6B). These results indicate a sharp differentiation between ‘bypass Sec14p’ mutants in their abilities to correct the derangements in nuclear division, normally associated with Sec14p dysfunction, even though these ‘bypass Sec14p’ mutations share in common their abilities to restore cell viability and efficient membrane trafficking to cells ablated for Sec14p function. These data also indicate that defects in the CDP-choline pathway of PtdCho synthesis do not significantly impact nuclear division.

**FIGURE 6.**
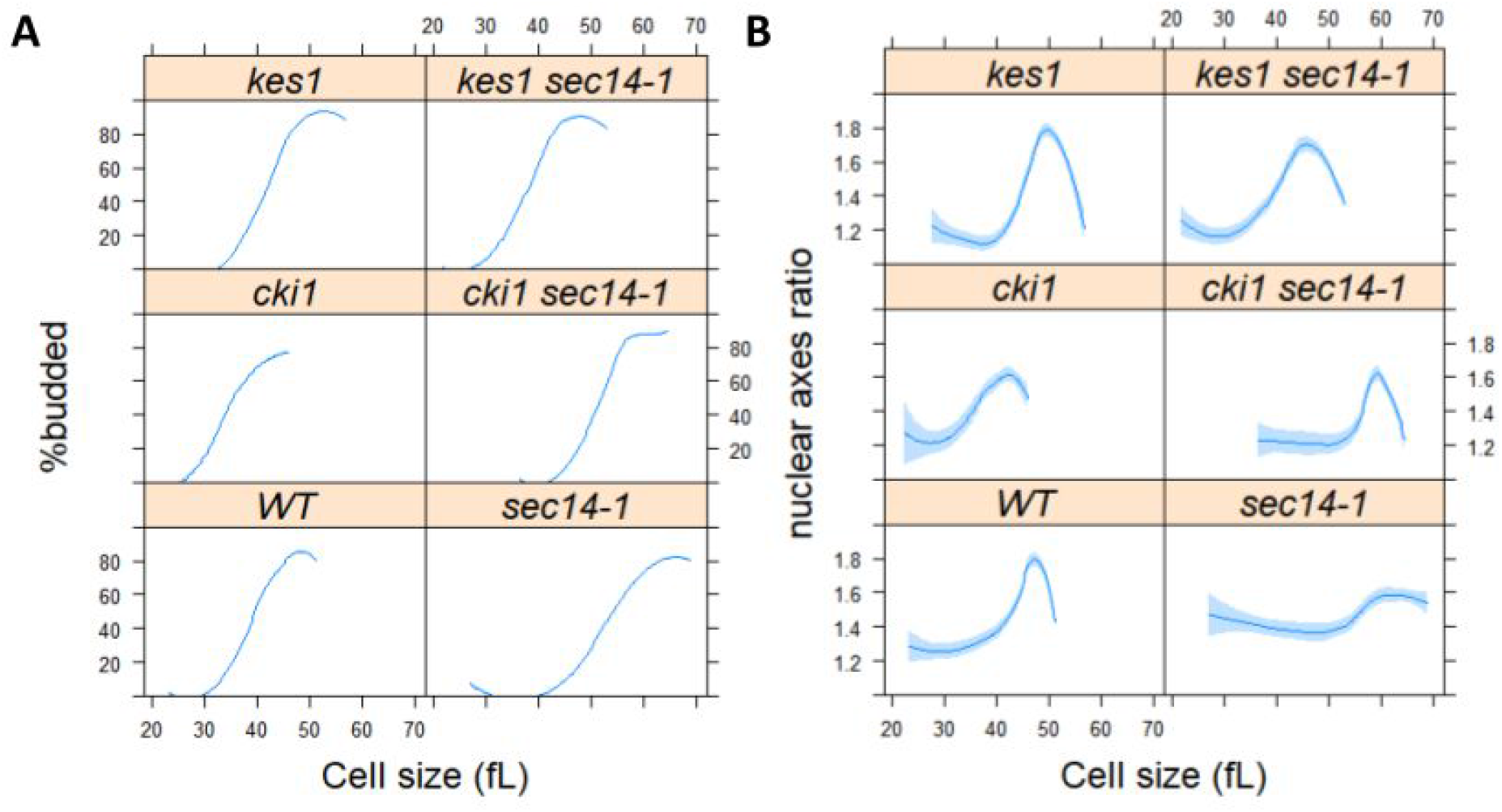
Loss of Keslp, but not Ckilp, suppresses the delayed nuclear division of *sec14-1* mutants. From synchronous cultures of the indicated strains, the percentage of budded cells (A) and nuclear shape (B) as a function of cell size (x-axis) were measured and plotted as in Figure 5. For the wild type cells (WT) and *sec14-1* cells, the values were the same as those in Figure 5. The values used to generate the graphs are in file Figure 6 - source data 1.

### Enhanced *ACC1* and *FAS1* translation corrects the aberrant nuclear shape of PtdIns 4-OH kinase mutants

A major function of Sec14p in cells is to stimulate the activities of the Golgi/endosomal and plasma membrane PtdIns 4-OH kinases Pik1p and Stt4p, respectively (Schaaf et al., 2008; Bankaitis et al., 2010). We therefore interrogated the cell cycle and nuclear morphogenetic phenotypes of temperature-sensitive mutants defective in the activities of these two lipid kinases as each enzyme is individually essential for yeast cell viability and these two enzymes together account for nearly all of the PtdIns 4-OH kinase activity in cells (Audhya et al., 2000). Even at their permissive temperature (25°C), *pik1-101* and *stt4-4* cells grew slowly and exhibited aberrant cell morphologies. Indeed, we were unable to obtain synchronous cultures of these mutants by elutriation. In an alternative approach, asynchronous cultures of *pik1-101* and *stt4-4* cells were examined for cell cycle parameters including cell size and DNA content. At the permissive temperature of 25°C, both *pik1-101* and *stt4-4* cells were significantly larger than wild type cells (~50% and 25%, respectively; Figure 7 - supplementary figure 1, top). Moreover, the DNA contents of both mutants were also irregular (Figure 7 - supplementary figure 1, bottom). Although Pik1p inactivation results in a cytokinesis defect at 37°C (Garcia-Bustos et al., 1994), we did not obtain evidence of increased ploidy at 25°C. Rather, the DNA content profiles for both mutants consistently reported a substantial delay in passage through S phase (Figure 7 - supplementary figure 1, bottom).

The cell cycle phenotypes described above, when coupled with prior evidence that phospholipid and triacylglycerol synthesis impacts nuclear morphology (Barbosa et al., 2019; Siniossoglou, 2013; Webster et al., 2010), prompted us to examine the nuclear morphologies of *pik1-101* and *stt4-4* cells. These mutants exhibited morphologically deranged nuclei characterized by highly irregular shapes that were in stark contrast to the normal spherical shape (Figure 7). Remarkably, *uORFm-ACC1,FAS1* restored normal nuclear morphology to *pik1-101* mutants and, to a lesser extent, to *stt4-4* mutants (Figure 7). Although the effect did not reach statistical significance in the *stt4-4* context (p=0.08, based on the Kruskal Wallis and posthoc Nemenyi tests; see also Figure 7 - figure supplement 2), the *pik1-101* data indicated that increased lipogenesis levied a significant correction of the abnormal nuclear shape of PtdIns 4-OH kinase mutants. These results further support the notion that lipogenesis actively controls nuclear shape and division, and identify a role for PtdIns-4-phosphate signaling in homeostatic control of nuclear morphology.

**FIGURE 7.**
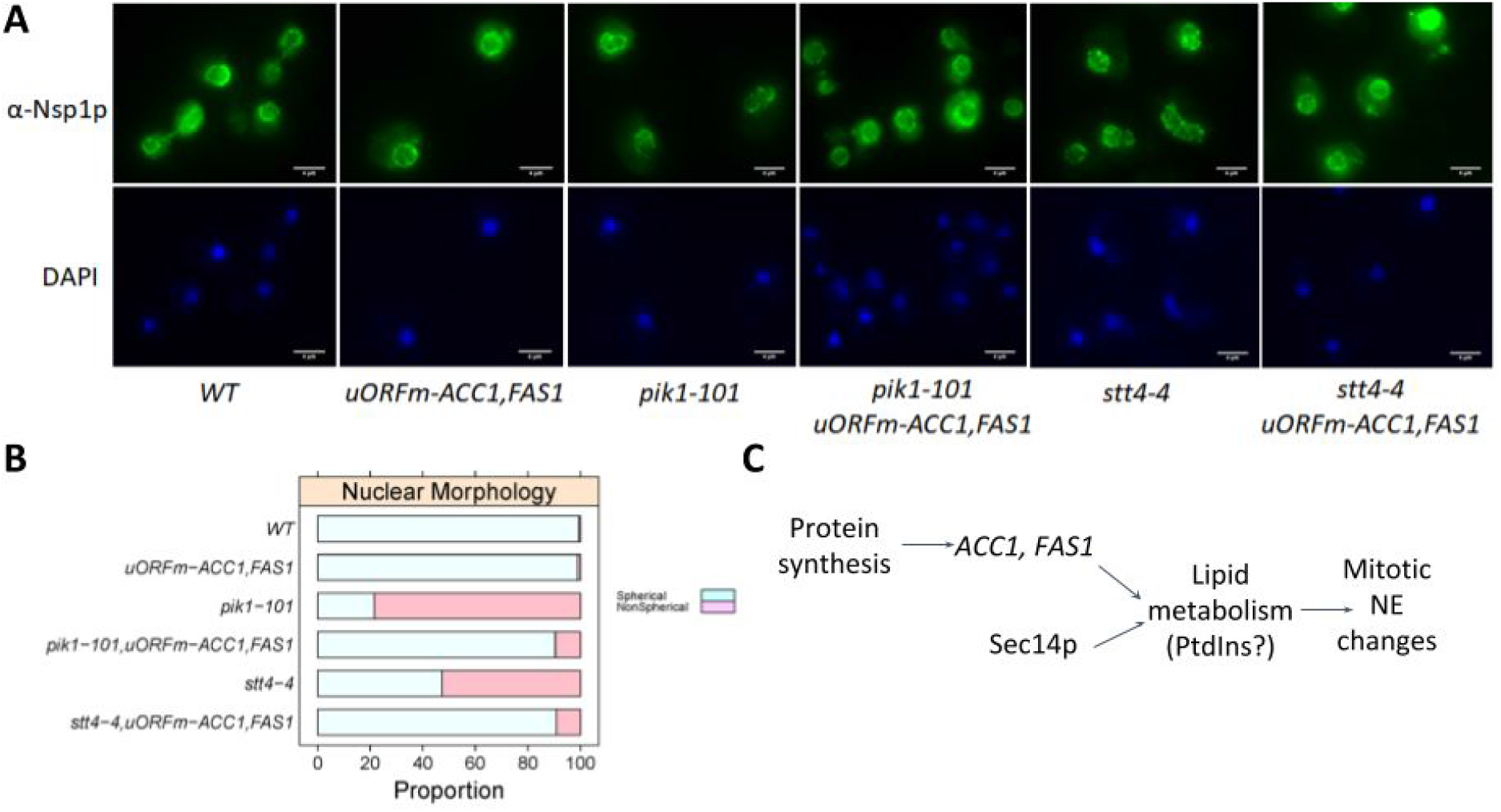
Upregulating translation of *ACC1* and *FAS1* corrects the aberrant nuclear shape of PI- 4-kinase mutants. A, Representative fluorescent microscopy images of cells from the indicated strains, stained with an α-Nsp1 antibody (top panels) or DAPI (bottom panels). B, Summary of the proportion of cells with spherical, or not, nuclei from the strains shown, calculated from three independent experiments (see FIGURE 7 - figure supplement 1). C, Schematic summary of our results.

## DISCUSSION

Herein, we demonstrate that gain-of-function mutations affecting translational control of lipogenic enzymes accelerate nuclear division, a critical cell cycle process. These results provide the first example of a metabolic pathway that is not merely required for a cell cycle process but actively drives the process forward. These findings are discussed in the context of cell cycle-dependent translational control of lipogenesis as it relates to regulation of nuclear morphogenesis.

### Translational control of lipogenic enzymes

The question of how mammalian acetyl-CoA carboxylase (ACC1, ACACA) and fatty acid synthase (FASN) are regulated is of intense interest because of the central role these enzymes play in the control of food intake, their contributions to obesity, and their association with cancer (Kuhajda et al., 1994; Baenke et al., 2013; Beloribi-Djefaflia et al., 2016). ACC1 expression is increased in conjunction with FASN to upregulate lipid synthesis in human tumors. Multiple mechanisms have been associated with the levels and activity of these enzymes in mammals. The activities of these enzymes in mammals are subject to multiple layers of regulation, and the control of Acc1 activity by phosphorylation has garnered considerable attention (Kim, 1997). Acc1 is inactivated upon phosphorylation by AMPK and, as leptin activates AMPK, this regulation is a crucial aspect of the mechanism by which leptin suppresses appetite (Gao et al., 2007). Moreover, the AMPK-mediated inhibition of Acc1 is relieved just before cytokinesis, and this timing correlates with increased Acc1 enzymatic activity late in the Hela cell cycle (Scaglia et al., 2014). However, the precise role of phosphorylation in the cell cycle regulation of Acc1p remains unclear. Large scale phosphoproteomic analyses identify several phosphorylation sites on yeast Acc1p. The major site is at residue S1157, which corresponds to S1216 in the human enzyme (Hunkeler et al., 2016). Although this site is conserved in the Acc1p enzymes of all eukaryotes, there are no reports that Acc1p^S1216^ phosphorylation is of any regulatory consequence for mammalian Acc1p. Moreover, there is no evidence that Acc1p phosphorylation is subject to cell cycle control in yeast.

Our data identify translation control as a mechanism for regulating lipogenesis as a function of the cell cycle. We had previously shown that translation of the mRNAs encoding Acc1p and Fas1,2p peaks late in the cell cycle ((Blank et al., 2017b); see also Figure 1 - figure supplement 2). That this represents a broadly conserved layer of regulation is indicated by subsequent studies that showed translational control of human *ACC1* is required for the progression of quiescent T cells into a proliferative state (Ricciardi et al., 2018). These prior studies linking elevated Acc1p and Fas1p activity with increased cell proliferation established these enzymes are necessary for lipogenesis and cell division. In that regard, there are numerous other examples where metabolic and cell growth processes are required for various cell cycle events (Johnston et al., 1977).

The novelty of the current study rests on our demonstration that upregulating Acc1p and Fas1p (by abrogating translational inhibitory elements) is sufficient to drive a critical cell cycle process – i.e., nuclear division. Functional ablation of the inhibitory uORFs that lie upstream of each of the corresponding structural genes elevated the overall levels of these enzymes while preserving the cell cycle-dependent oscillations in their abundance, albeit with a dampened amplitude ((Blank et al., 2017b); see also Figure 1 - figure supplement 2). This is an expected result because, for the uORF-mediated control to fully account for the mitotic peak of translation of *ACC1* and *FAS1*, the cellular ribosome content must also fluctuate as cells progress through the cell cycle. Evidence from our lab and others argues against such cell cycle fluctuations in ribosome content (Blank et al., 2020; Elliott et al., 1979; Shulman et al., 1973). Our studies now pose the open question of what mechanisms contribute to the mitotic peak in *ACC1* and *FAS1 t*ranslation? This level of regulation is of clear physiological relevance. Whole-cell metabolic profiling demonstrates lipid abundance (including all major classes; PtdIns, PtdCho, PtdEtn, and triglycerides) is the most periodic metabolic parameter as a function of the cell cycle and that it peaks in mitosis (Blank et al., 2020). Furthermore, we now show that enhanced *ACC1* and *FAS1* translation was sufficient to increase most lipids’ abundances (Figure 2).

### Nuclear morphology and lipogenesis

Yeast and other fungi undergo closed mitosis, where the nuclear envelope remains intact throughout the entirety of the cell cycle. The nucleus maintains a spherical shape throughout interphase and then elongates during mitosis (Meseroll and Cohen-Fix, 2016).The fact that nuclear envelope expansion depends on cell cycle progression, and not on a general increase in cell size, argues for specialized metabolic inputs, including fatty acid and phospholipid synthesis (Walters et al., 2018). Indeed, nuclear membrane proliferation is required for progression through mitosis, regardless of whether it is closed or disassembled before reassembling during telophase as in mammalian cells. Inhibition of fatty acid synthase impairs nuclear elongation and chromosome segregation in fission yeast (Takemoto et al., 2016). A mitotic delay is occasionally accompanied by the appearance of nuclear membrane extensions, or flares, in budding yeast. These flares are interpreted as evidence for continued nuclear elongation on the face of some cell cycle arrest (Walters et al., 2012; Meseroll and Cohen-Fix, 2016). In support of this interpretation, *ACC1*, *FAS1, or FAS2* loss-of-function mutations impair flare formation and nuclear elongation (Walters et al., 2014). However, we emphasize that the acceleration of nuclear elongation and division that we observed in the face of the upregulated translation of lipogenic enzymes is without precedent. Those data are not only fully consistent with the requirement lipogenesis in nuclear elongation and division, but these also establish that enhanced lipogenesis is sufficient to drive these processes.

### Phosphoinositide signaling in control of nuclear division and morphology

While the role of the normally essential Sec14p in membrane trafficking is well documented (Bankaitis et al., 1989; Cleves et al., 1989; Fang et al., 1996), there is evidence that the essential cellular function(s) executed by Sec14p are not solely at the level of membrane trafficking control but also at the level of cell cycle control (Mousley et al., 2012). In that regard, Sec14p hypomorphisms manifest themselves in cell cycle phenotypes such as delay in the passage through G2/M and progression from G1 into S-phase (Huang et al., 2018). In that context, we report an unexpected confluence of translational control of lipogenesis with Sec14p-dependent phosphoinositide signaling in the cell cycle. We find that *sec14-1* mutants are not only delayed in the initiation and completion of nuclear division, but that gain-of-function mutations that functionally ablate the *ACC1* and *FAS1* uORFs (with the result that the two corresponding mRNAs are translationally derepressed) partially rescue *sec14-1* growth defects. Moreover, functional ablation of the uORFs fully rescued the delays in progression through G2/M and G1/S in Sec14p-deficient mutants. What is particularly striking in light of those data is that the accelerated timing of nuclear division in the uORF mutants was corrected in the *sec14-1ts* genetic background. These collective data not only indicate that increased lipogenesis suppresses the mitotic functions of Sec14p but also suggest that the accelerated nuclear division schedule associated with elevated lipogenesis requires Sec14p involvement.

The available evidence indicates that the Sec14p involvement was independent of its role in membrane trafficking on two counts. First, these phenotypes were manifested at permissive temperatures for *sec14-1* mutants where membrane trafficking pathways are operating normally by all measurable criteria. Second, the ‘bypass Sec14’ mutant that inactivates Cki1p (i.e., the choline kinase representing the first step in the CDP-choline pathway for PtdCho biosynthesis) corrects the membrane trafficking defects of *sec14* mutants but fails to correct the deranged scheduling of nuclear elongation and division in *sec14-1* mutants. Rather, as Sec14p stimulates PtdIns-4-P synthesis by both the Pik1p and Stt4p kinases in vivo, the data argue that Sec14p-mediated phosphoinositide signaling is involved in regulating nuclear elongation and division. Two lines of evidence support this interpretation of the data. First, inactivation of Kes1p, an antagonist of Sec14p- and Pik1p-dependent PtdIns-4-P signaling (Fang et al., 1996; Li et al., 2002), corrects both the membrane trafficking and the nuclear elongation/division defects associated with Sec14p deficiencies. In this manner, the *kes1* ‘bypass Sec14p’ mutants differ from *cki1* mutants that correct only the membrane trafficking deficits. Second, cells compromised for Pik1p activity presented misshapen nuclei whose morphological derangements were corrected by increased lipogenesis. These data are intriguing in light of previous demonstrations that Pik1p shuttles between cytosolic and nuclear pools and executes essential cellular functions in both compartments (Garcia-Bustos et al., 1994; Strahl et al., 2005).

Precisely how PtdIns-4-P signaling modulates nuclear division remains to be determined. One execution point could be at the translational control level as inactivation of Pik1p or Stt4p triggers a block in protein synthesis through phosphorylation of translation initiation factor eIF2α (Cameroni et al., 2006). Another attractive and not mutually exclusive possibility is that PtdIns-4-P serves as a precursor to synthesizing a nuclear pool of PtdIns-4,5-bisphosphate (PtdIns-4,5-P2). There is abundant literature regarding nuclear PtdIns-4,5-P2 and its roles (direct and indirect) in regulating multiple nuclear events such as transcription, mRNA processing and polyadenylation, and mRNA export (Cocco et al., 1987, 1989; York et al., 1999; Li et al., 2013; Hamann and Blind, 2018; Chen et al., 2020). We attempted to assess the relationship between PtdIns-4,5-P2 synthesis and *ACC1* and *FAS1* translational control but were unsuccessful in generating the requisite strains for the analysis. So, the relationship between PtdIns-4-P and PtdIns-4,5-P2 signaling related to nuclear elongation and division remains to be resolved.

In summary, the results reported herein emphasize the physiological significance of generating specific lipid pools that drive a crucial cell cycle process – i.e., nuclear elongation and division. These lipids pools represent metabolic inputs that do not merely ‘fuel’ a cell cycle process but, rather, play heretofore unappreciated active and instructive signaling roles in driving landmark cell cycle events. The collective data integrate protein synthesis, lipogenesis and phosphoinositide signaling with the timing of nuclear elongation and division. In this manner, this study sets the stage for deciphering the specific identities and compartmental organization of the relevant lipid pools that control these signaling processes.

## MATERIALS AND METHODS

### Key resources table

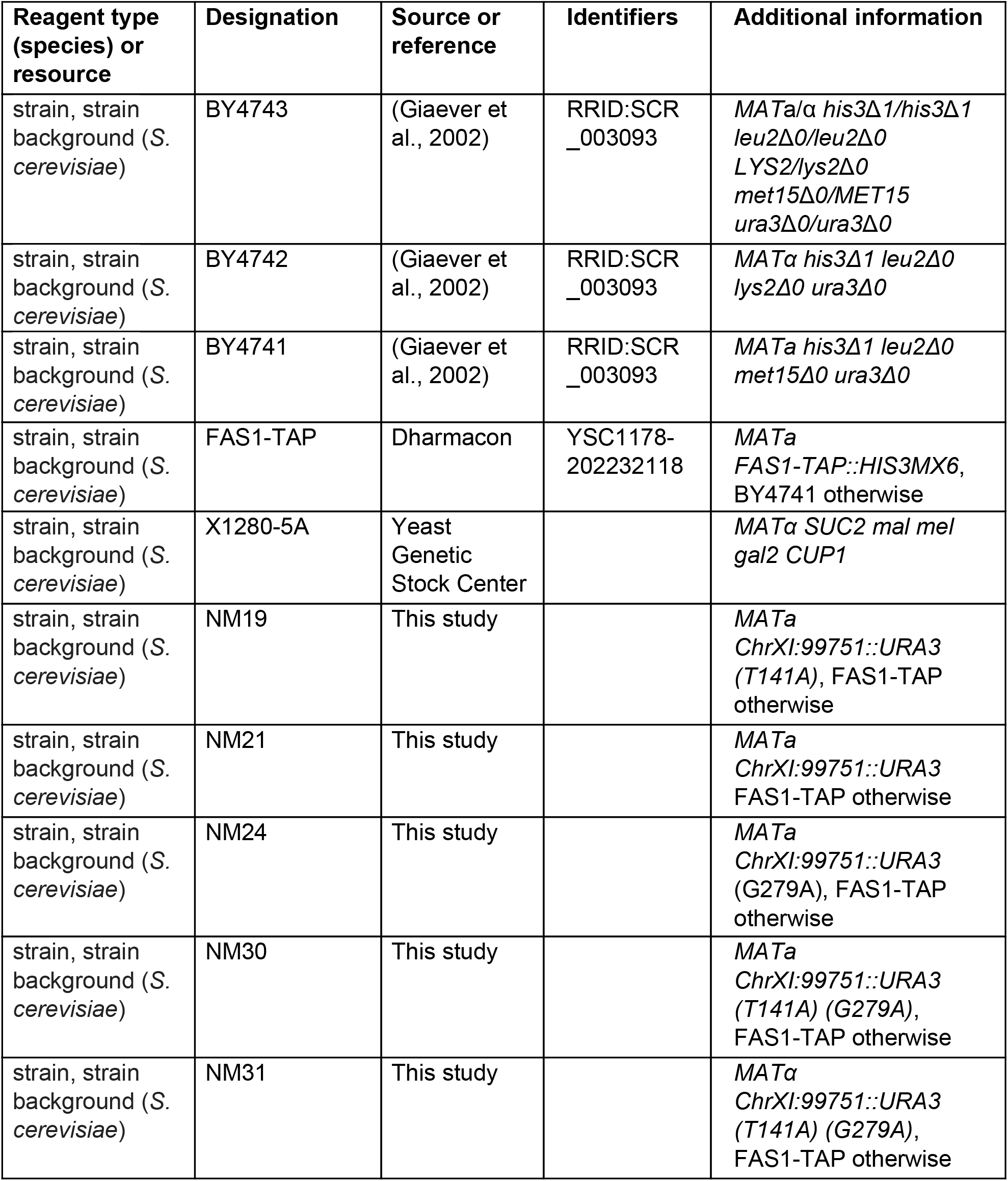
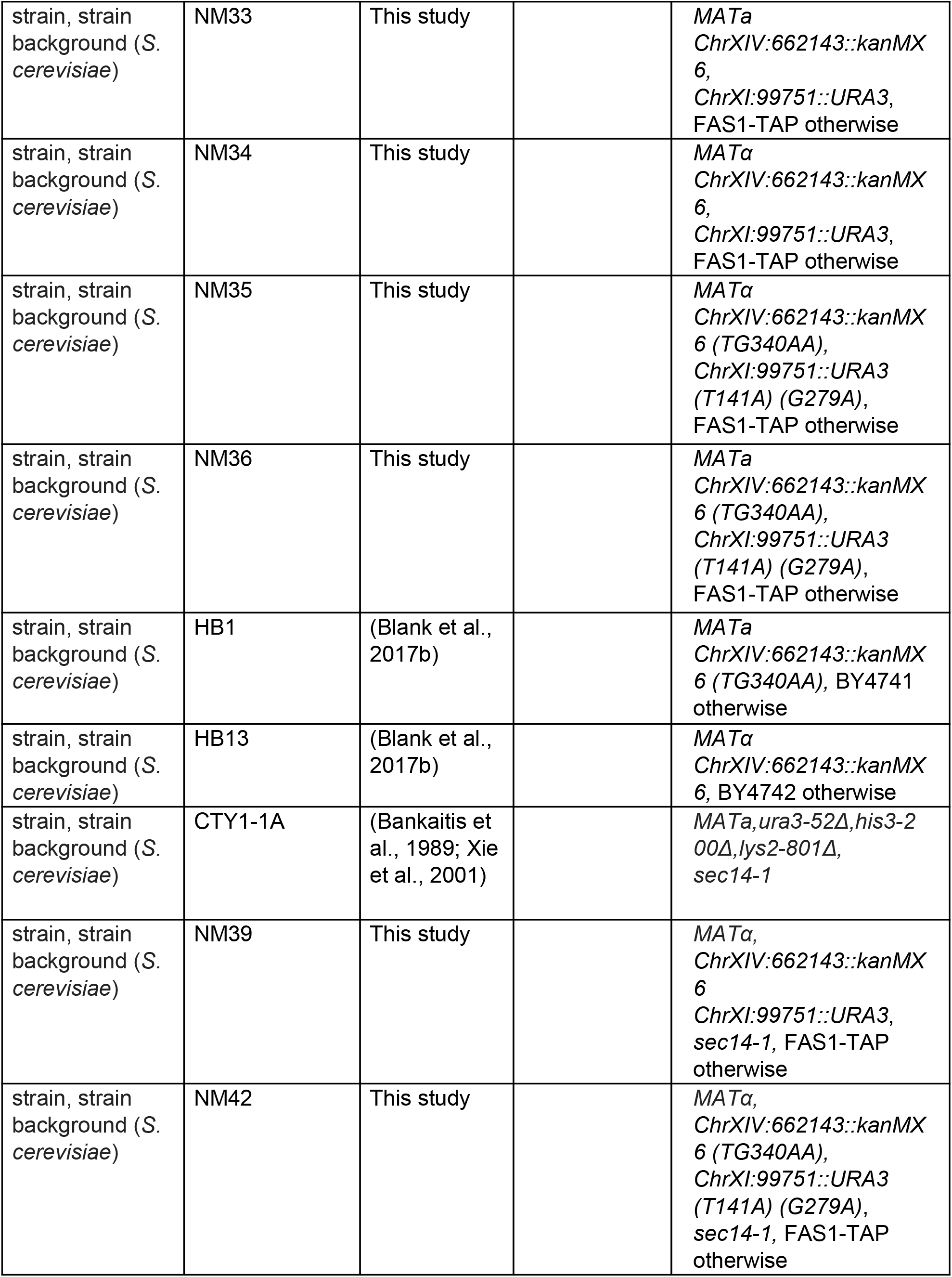
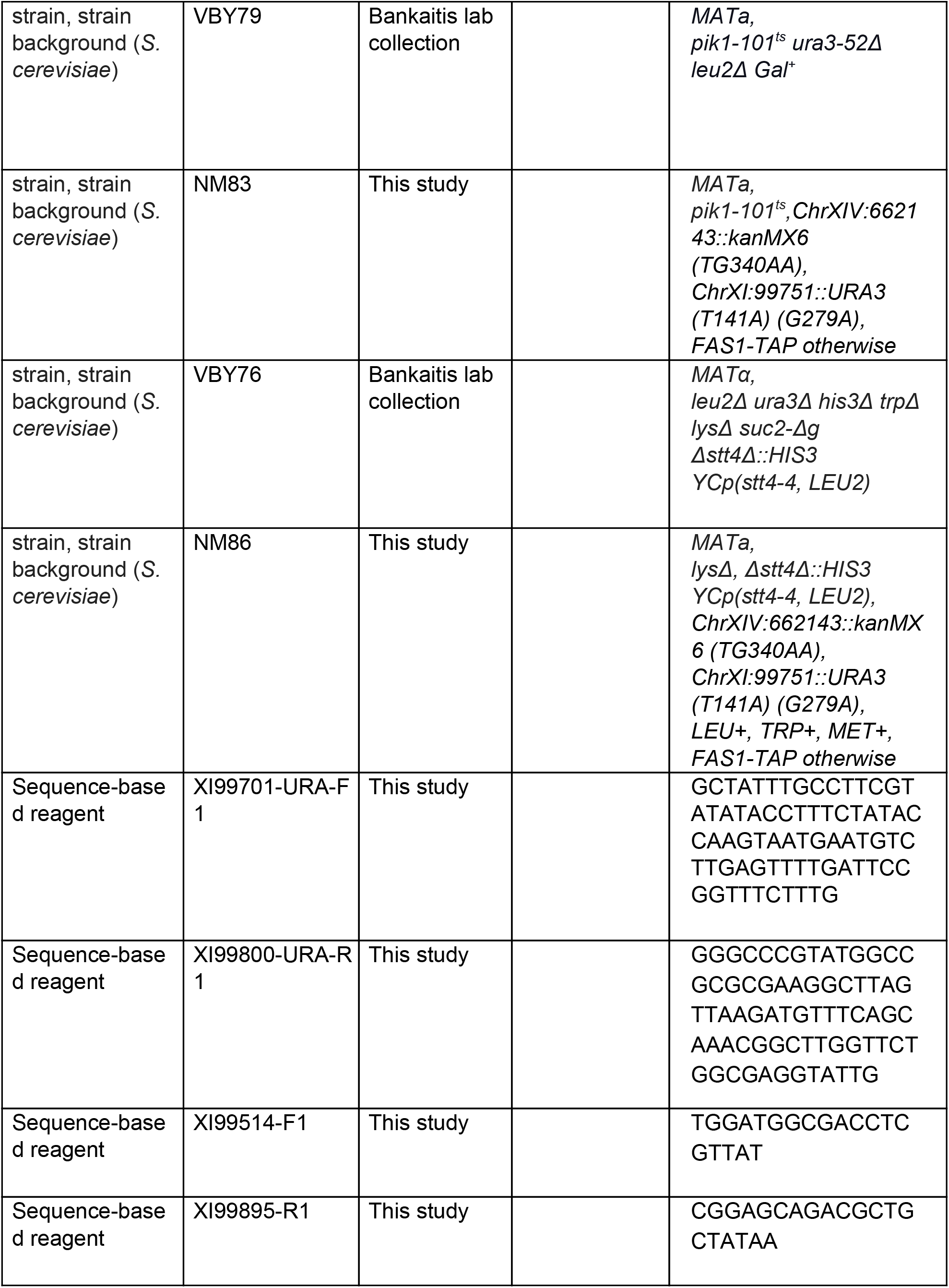
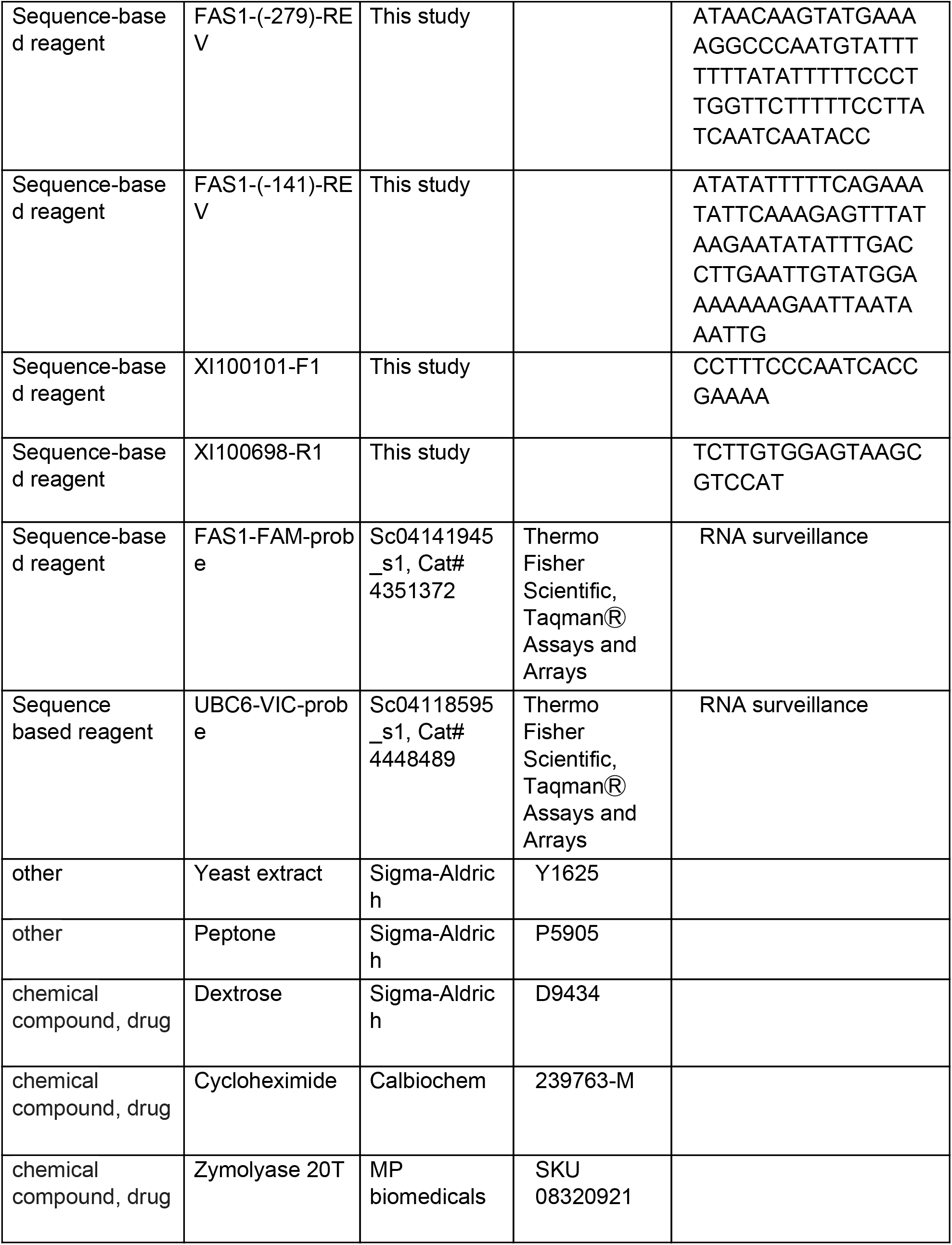
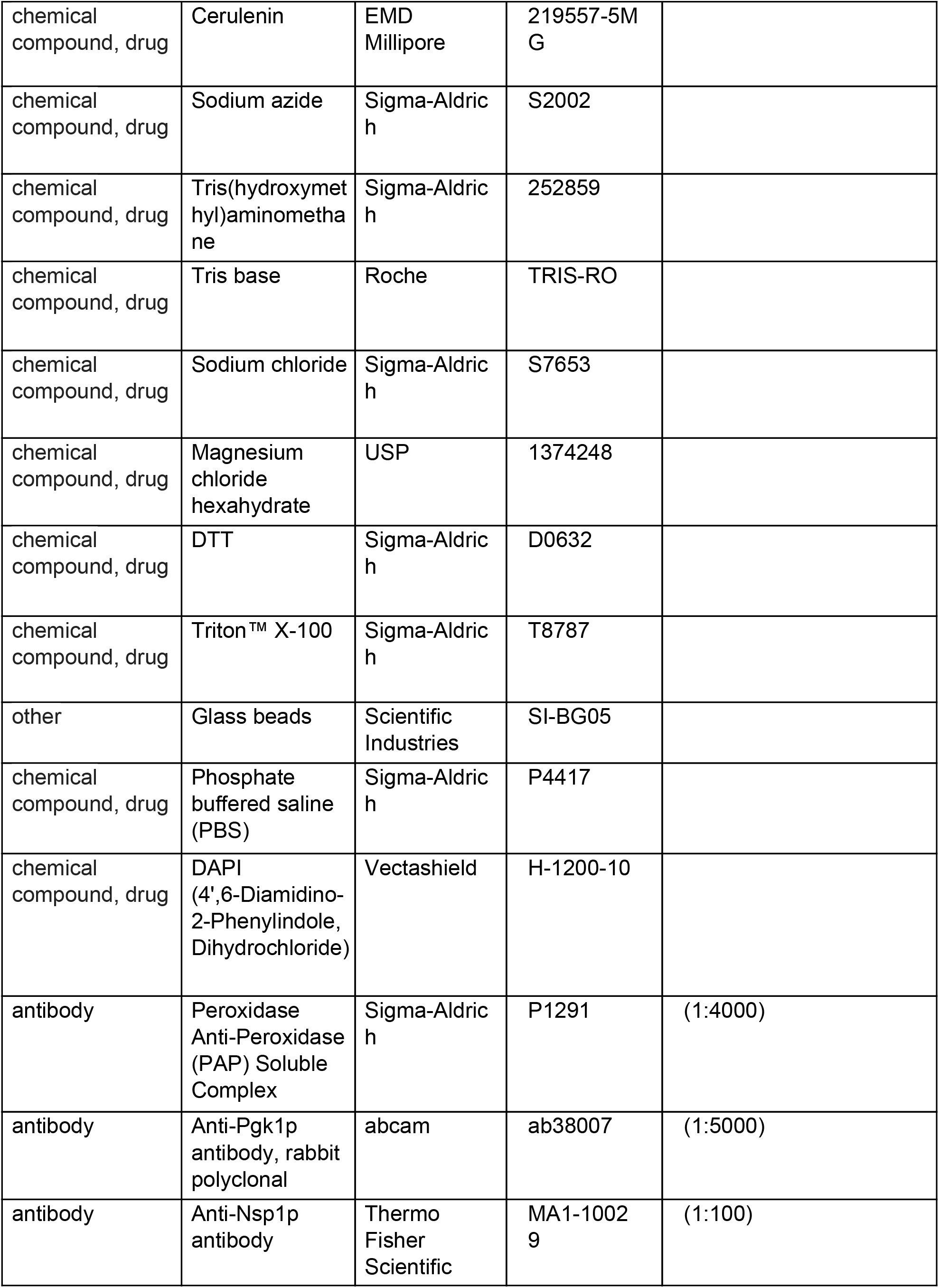
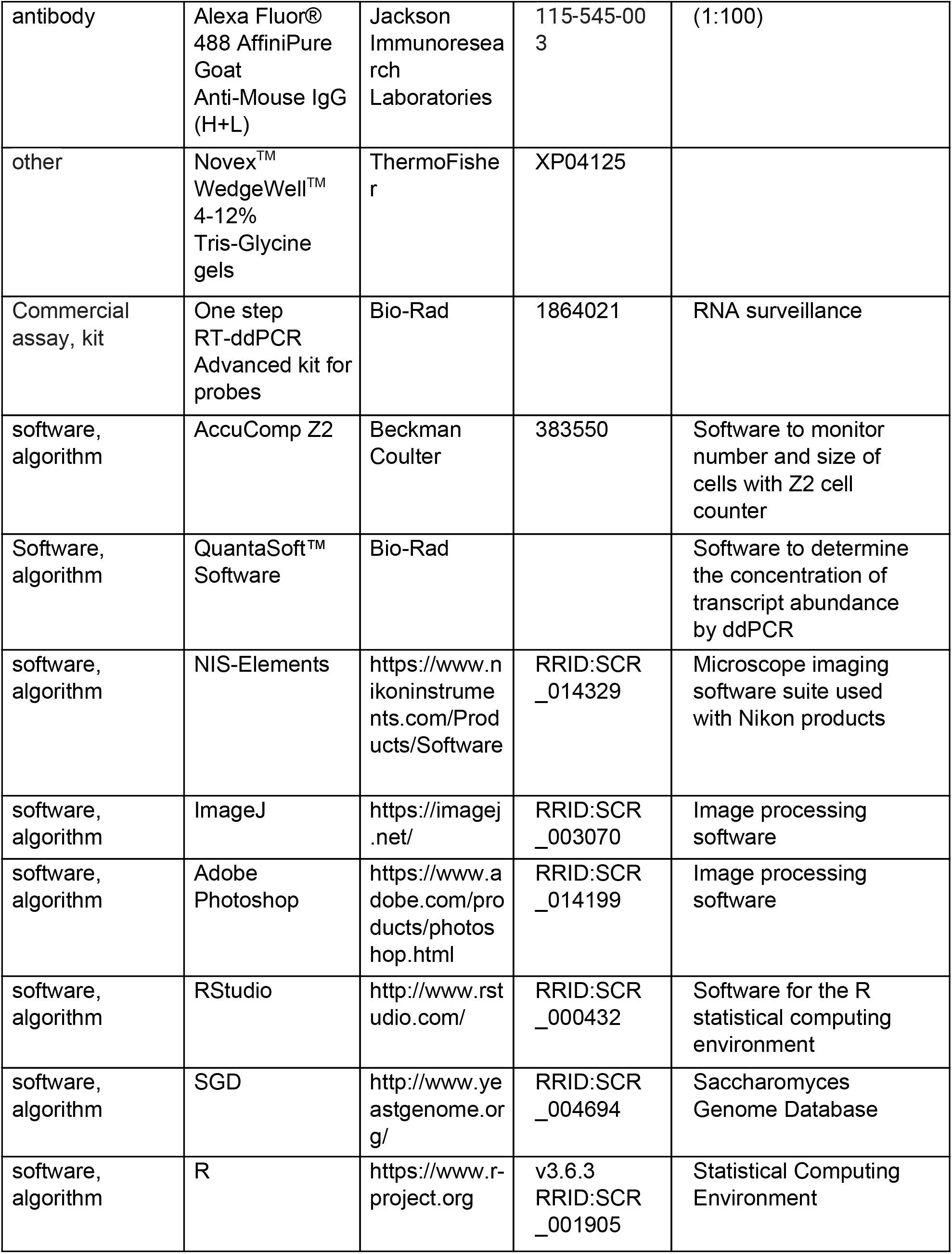

Where known, the Research Resource Identifiers (RRIDs) are shown.

### Strains and media

All the strains used in this study are shown in the Key Resources Table, above. Unless noted otherwise, the cells were cultivated in the standard, rich, undefined medium YPD (1% ^w^/_v_ yeast extract, 2% ^w^/_v_ peptone, 2% ^w^/_v_ dextrose), at 30°C (Kaiser et al., 1994). To modify the *FAS1-5’* leader, we first inserted a *URA3* gene (amplified from a prototrophic strain X1280) upstream, at position ChrXI99751 with the PCR-mediated methodology of (Longtine et al., 1998), using primers XI99701-URA-F1 and XI99800-URA-R1 (see Key Resources Table). The PCR product was transformed into a yeast strain carrying the TAP epitope at the 5’ end of the *FAS1* main ORF (FAS1-TAP, see Key Resources Table). The resulting strain, NM21, was genotyped by PCR for the presence of the insertion at the expected location, using primers (XI99514-F1 and XI99895-R1) that flank the insertion. Genomic DNA of NM21 was then used as a template in a PCR reaction with the forward primer XI99514-F1 and reverse primers FAS1-141-REV, or FAS1-279-REV, to introduce point mutations at the start codon of the proximal uORF (T141A), or the distal uORF (G279A), respectively. The strains obtained were mutant for the proximal (NM19), or distal uORF (NM24). The introduced mutations were verified by PCR (with primers XI100101-F1 and XI100698-R1) followed by restriction digestion with enzymes PsiI (for the proximal uORF mutation) and RsaI (for the distal uORF mutation). The T141A mutation introduces a restriction site for PsiI at the proximal uORF. On the other hand, the G279A mutation eliminated the RsaI restriction site at the distal uORF. Next, to obtain a double uORF *FAS1* mutant, the genomic DNA of NM24 was used as a template in a PCR reaction with forward primer XI99514-F1, and reverse primer FAS1-141-REV. The resulting product was transformed into FAS1-TAP cells, to generate strain NM31, which was verified by PCR, followed by restriction digestion PsiI. The introduced uORF mutations were also confirmed by sequencing the strains’ genomic DNA using the primers XI100101-F1 and XI100698-R1.

To de-repress translation of both *ACC1* and *FAS1* in the same cells, we crossed strain NM31 with the *ACC1-uORF* mutant (carrying the TG340AA substitutions in the 5’-leader of *ACC1*; strain HB1, see Key Resources Table and (Blank et al., 2017b)), followed by sporulation and tetrad dissection. The resulting strains (NM36, NM35) have point mutations at the start codons of the uORFs of both *ACC1* and *FAS1* (*uORFm-ACC1, FAS1*). To obtain wild type strains with markers matching those of *uORFm-ACC1, FAS1* cells, we crossed NM21 with HB13 (ChrXIV 662143::kanMX6, see (Blank et al., 2017b)), followed by sporulation and tetrad dissection, yielding strains NM33 and NM34. Unless indicated otherwise, strains NM33, and NM36, were used as the wild type, and *uORFm-ACC1, FAS1*, strains shown in Figures.

To introduce the *sec14-1* allele, strains NM34 and NM35 were crossed with CTY1-1A (Xie et al., 2001), sporulated, and the tetrads dissected to generate the marker-matched *sec14-1* (NM39), and the *sec14-1, uORFm-ACC1, FAS1* quadruple mutant (NM42), respectively. Similarly, we generated all other mutants combinations shown in Figures 5, 6.

### Sample-size and replicates

For sample-size estimation, no explicit power analysis was used. All the replicates in every experiment shown were biological ones, from independent cultures. A minimum of three biological replicates was analyzed in each case, as indicated in each corresponding figure’s legends. Where three replicates were the minimum number of samples analyzed, the robust bootstrap ANOVA was used to compare different populations via the t1waybt function, and the posthoc tests via the mcppb20 function, of the WRS2 R language package (Wilcox, 2011; Mair and Wilcox, 2016). For measurements where at least four independent replicates were analyzed, we used non-parametric statistical methods, as indicated in each case. No data or outliers were excluded from any analysis.

### Immunoblot analysis

For protein surveillance, protein extracts were made as described previously (Amberg et al., 2006), and run on 4–12% Tris-Glycine SDS-PAGE gels. To detect TAP-tagged proteins with the PAP reagent, we used immunoblots from extracts of the indicated strains, as we described previously (Blank et al., 2017b, 2020). Loading was evaluated with an anti-Pgk1p antibody. Images were processed with the ImageJ software package. Briefly, the ‘Subtract background’ tool was applied, followed by the ‘Measure’ tool to obtain a mean intensity for each band. The area measured was kept constant for a sample series for each blot analyzed.

### Transcript abundance using ddPCR

For RNA surveillance, RNA extracts were made as described previously (Blank et al., 2017b). Briefly, RNA was extracted using the hot acidic phenol method (Collart and Oliviero, 2001). Cell pellets were resuspended in 0.4 ml of TES buffer (10 mM Tris, pH 7.5; 10 mM EDTA, and 0.5% SDS) with 0.1 ml of glass beads. Then, 0.4 ml of acid phenol, pH 4.5, was added, and the samples were heated at 65°C for 0.5 h with occasional vortexing for 10 s each time. The samples were then centrifuged for 5 m at 13,000 *g*. The aqueous layer was transferred to a new tube containing 1 ml of cold 100% ethanol with 40 μl of 3 M sodium acetate and incubated overnight at 4°C. The next day, the samples were centrifuged at 13,000 *g* for 20 min at 4°C, washed with 80% ethanol, and centrifuged again for 5 min at 13,000 *g*. The pellets were resuspended in 25 μl water. The amount of total RNA in each sample was measured with a spectrometer. For the quantification of transcript abundance, 0.75 ng of total RNA was used for each sample.

The ddPCR reaction mixture was prepared by following the manufacturer’s protocol, using the Taqman hydrolysis probes labeled with FAM (for *FAS1*) and VIC (for *UBC6*) reporter fluorophores. The mixture was kept on ice throughout the whole experiment. Once the reaction mixture was prepared, the samples were placed into a droplet generator (QX200™ AutoDG™ Droplet Digital™ PCR System), which uses specially developed reagent and microfluidics to partition each sample into 20,000 nl-sized droplets. Once the droplets were generated, the samples were transferred to a thermal cycler (C1000 Touch™ Thermal cycler) for PCR. Following the PCR, the plate containing the droplets was placed in a droplet reader (Bio-Rad, QX200 Droplet reader). The autosampler of the droplet reader picks up droplets from each of the wells of the PCR plate, and fluorescence is measured for individual droplets.

The abundance of the transcripts was obtained using the QuantaSoft™ Software. Transcript levels of *FAS1* were normalized against the corresponding transcript levels of *UBC6*.

### Centrifugal elutriation, cell size and DNA content measurements

All methods have been described previously (Hoose et al., 2012; Soma et al., 2014).

### Metabolite profiling

The untargeted, primary metabolite and complex lipid analyses were done at the NIH-funded West Coast Metabolomics Center at the University of California at Davis, according to their established mass spectrometry protocols, as we described previously (Blank et al., 2020; Maitra et al., 2020). The raw data for the complex lipid measurements are in file Figure 2 - source data 1. The raw data for the primary metabolite measurements are in file Figure 2 - source data 2. To identify significant differences in the comparisons among the different strains, we used the robust bootstrap ANOVA, as described above. The input values we used were first scaled-normalized for input intensities per sample. Detected species that could not be assigned to any compound were excluded from the analysis.

### Fluorescence microscopy

Cells were fixed with 3.7% ^v^/_v_ formaldehyde at room temperature for 30 m, washed three times with 0.1 M potassium phosphate buffer, followed by a wash and resuspension in 1.2 M sorbitol. Next, the fixed cells were digested with 0.06 mg/ml zymolyase 20T for 5 m at 37°C, and later washed and resuspended again in 1.2 M sorbitol. 20 μl of The fixed and digested cells were added on a poly-lysinated slide and incubated at room temperature in a moist chamber for 20 m. The liquid was removed, and the slide was dehydrated by ice-cold methanol, and then acetone, for 3 m, and 10 s, respectively. The slide was air-dried and incubated in 0.1 % BSA/PBS for 30 m at room temperature. The BSA/PBS solution was removed, and the α-Nsp1 antibody (See Key Resources Table) was added at a dilution of 1:100 dilution and incubated overnight at 4°C. Nsp1p is a nucleoporin (Hurt, 1988) used in our experiments to decorate the nuclear envelope (Figure 3 - figure supplement 1). The next day, the slide was washed five times with 0.1% BSA/PBS, followed by the addition of secondary antibody (Alexa Fluor 488 AffiniPure Goat α-Mouse IgG, See Key Resources Table) at a dilution of 1:100 and incubated at room temperature for 1 h. The slide was then washed with 0.1% BSA/PBS and mounted with the Vectashield mounting media containing DAPI to visualize the nuclear DNA.

Cells were viewed with a Nikon Eclipse TS100 microscope, with a 100X objective, and the images were captured with a CoolSnap Dyno 4.5 Nikon camera. The exposure time for the DAPI and GFP filters (Alexa Fluor 488) was 50 ms and 500 ms, respectively. All images were captured in NIS Elements Advanced Research (version 4.10) software. The fluorescent images acquired with the GFP and DAPI filters were processed in ImageJ. The ratio of two axes (long axis and the short axis) of the nuclear mass were measured using the ‘line’ tool followed by the ‘measure’ tool in the ImageJ software package. When the nucleus is near-spherical, the ratio is 1. However, as the nucleus starts to expand, the ratio increases, signifying the elongation of the nuclear envelope (Figure 3 - figure supplement 1).

## ACKNOWLEDGEMENTS

This work was supported by NIH grant R01GM123139 to M.P. V.A.B. was supported by NIH grant R35 GM131804.

---

**FIGURE 1 - figure supplement 1.**
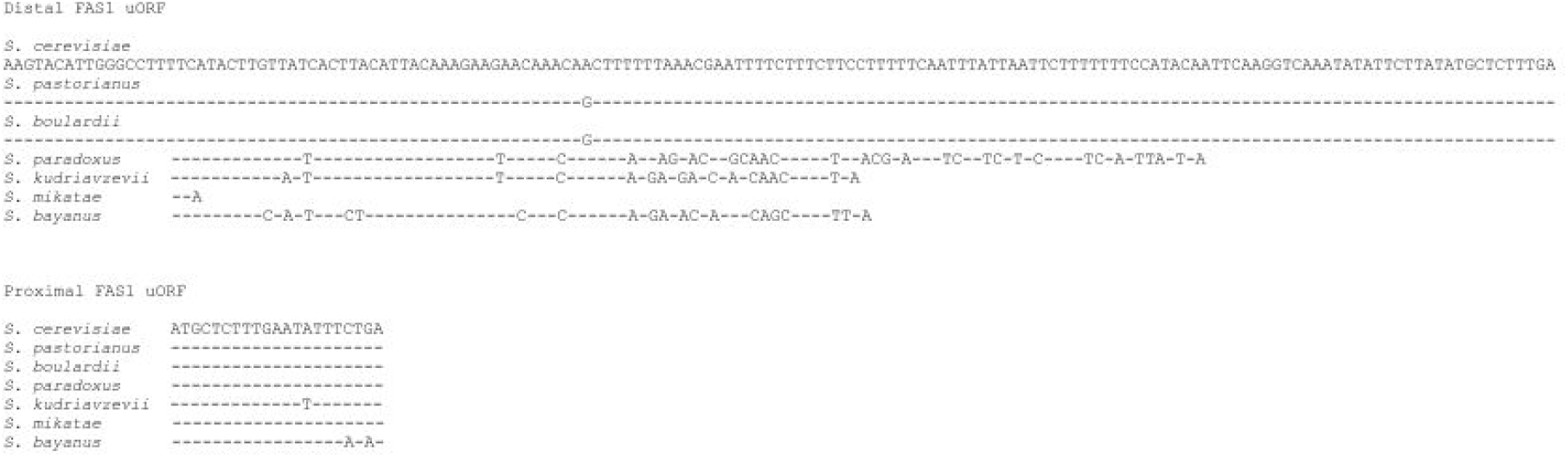
Multiple sequence alignment of the *FAS1* uORFs among Saccharomyces species. Sequences corresponding to the distal (Top) and proximal (Bottom) *FAS1* uORF sequences. The sequences were aligned based on the output of the Fungal BLAST function from the Saccharomyces Genome Database (SGD).

**FIGURE 1 - figure supplement 2.**
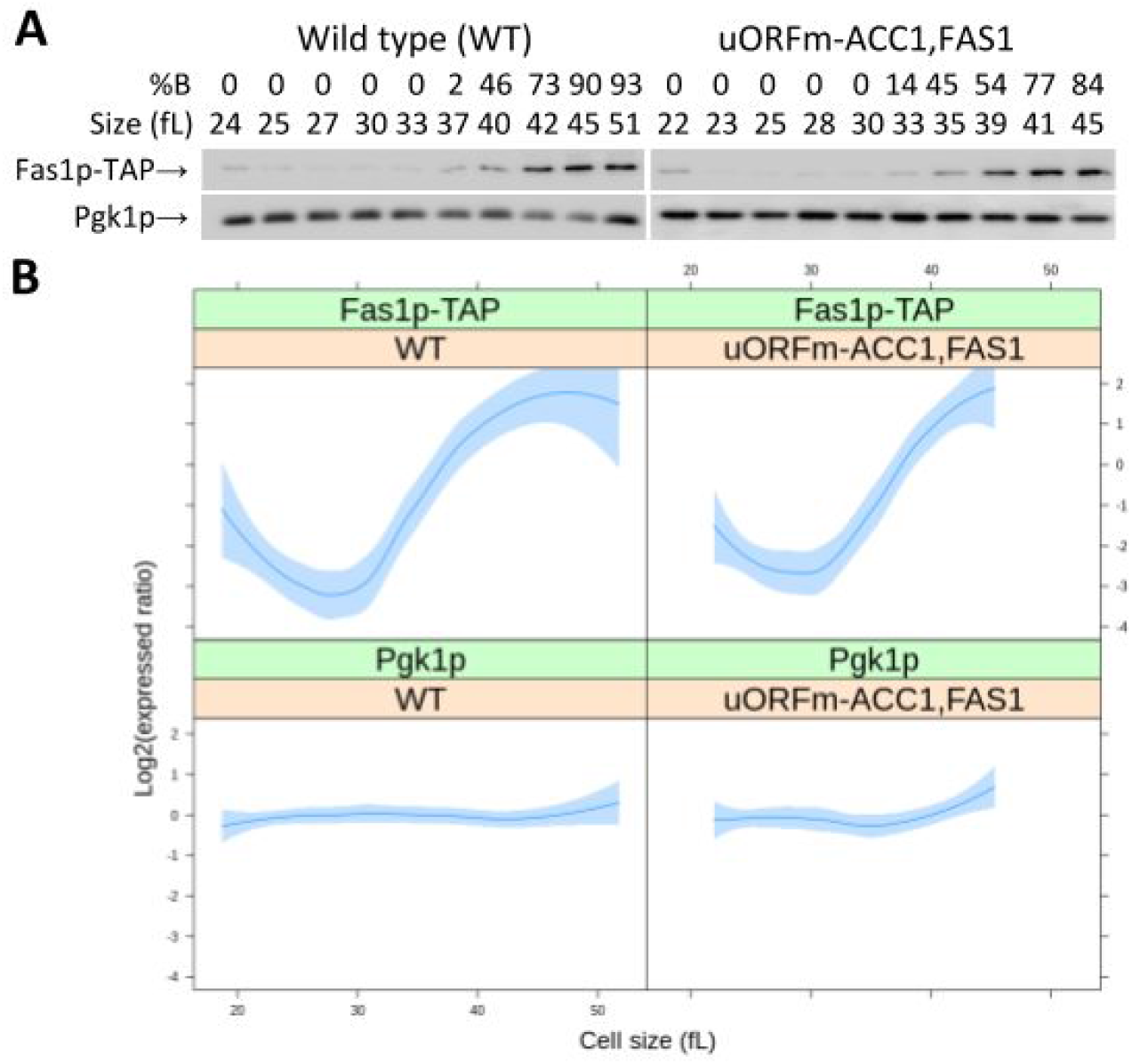
Fas1p abundance is periodic in cells lacking the uORFs in *ACC1* and *FAS1*. The abundance of Fas1p-TAP was monitored in wild type and *uORFm-ACC1, FAS1* cells, constructed as described in Materials and Methods. Samples were collected by elutriation in a rich, undefined medium (YPD) and allowed to progress synchronously in the cell cycle. Experiment-matched loading controls (measuring Pgk1p abundance) were also quantified and shown in parallel. A, Representative immunoblots, along with the percentage of budded cells (%B) and the cell size (in fL), for each of the samples. B, From at least three independent experiments in each case, the Fas1p-TAP and Pgk1p signal intensities were quantified as described in Materials and Methods. The Log2(expressed ratios) values are on the y-axis, and cell size values are on the x-axis. Loess curves and the standard errors at a 0.95 level are shown. All the immunoblots for this figure are in file Figure 1 - source data 3, while the values used to generate the graphs are in file Figure 1 - source data 4.

**FIGURE 2 - figure supplement 1.**
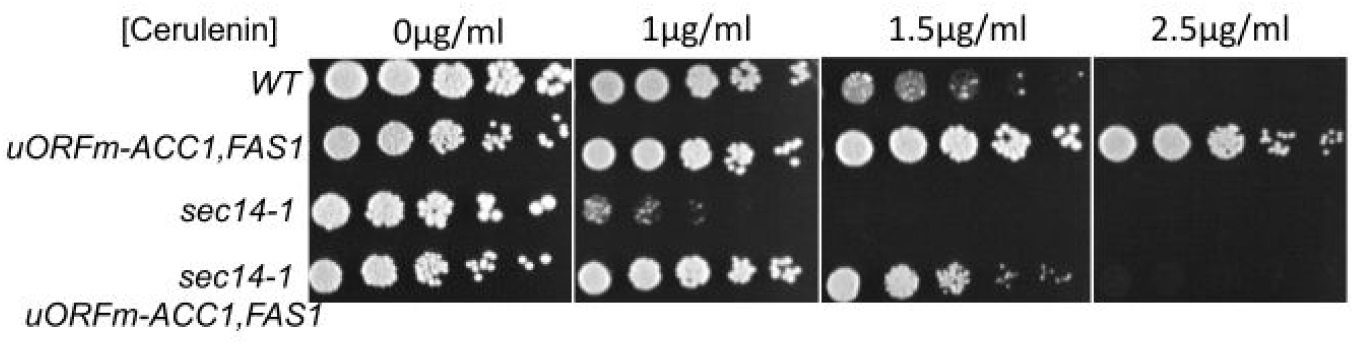
Cells lacking the uORFs in *ACC1* and *FAS1* are resistant to cerulenin, a specific FAS inhibitor. Serial dilutions (5-fold) of cultures of the indicated strains were spotted on solid media containing the indicated concentration of cerulenin, and photographed after 3 days.

**FIGURE 3 - figure supplement 1.**
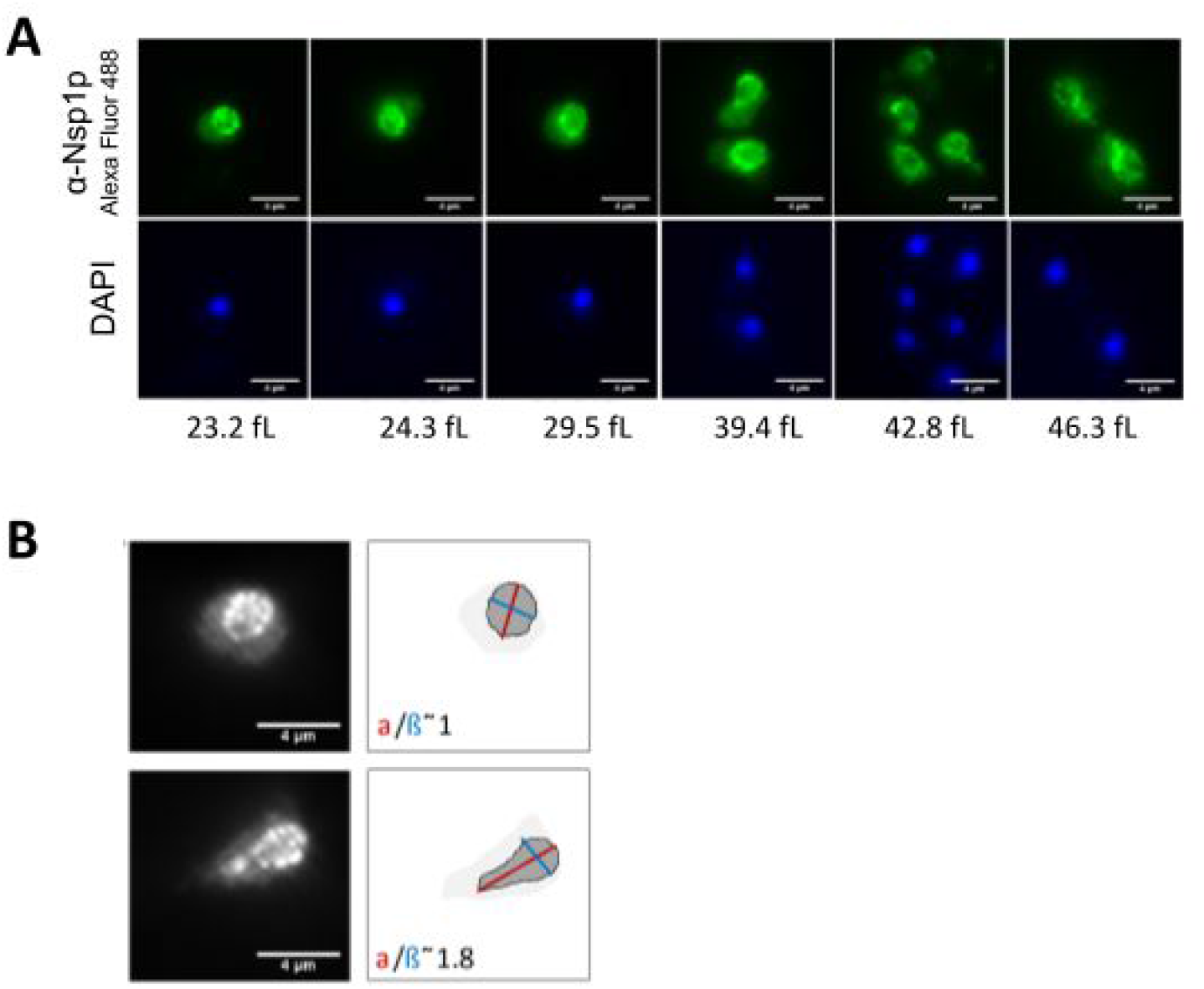
Monitoring nuclear morphology. A, Representative fluorescent microscopy images of elutriated, wild type cells, stained with an α-Nsp1 antibody (top panels) or DAPI (bottom panels). The mean cell size (in fL) of the samples are shown below the corresponding panels. B, Images of yeast cells, with the Nsp1p signal decorating spherical (top) and elongated (bottom) nuclei. The axes we used to score nuclear shape are on the schematic.

**FIGURE 7 - figure supplement 1.**
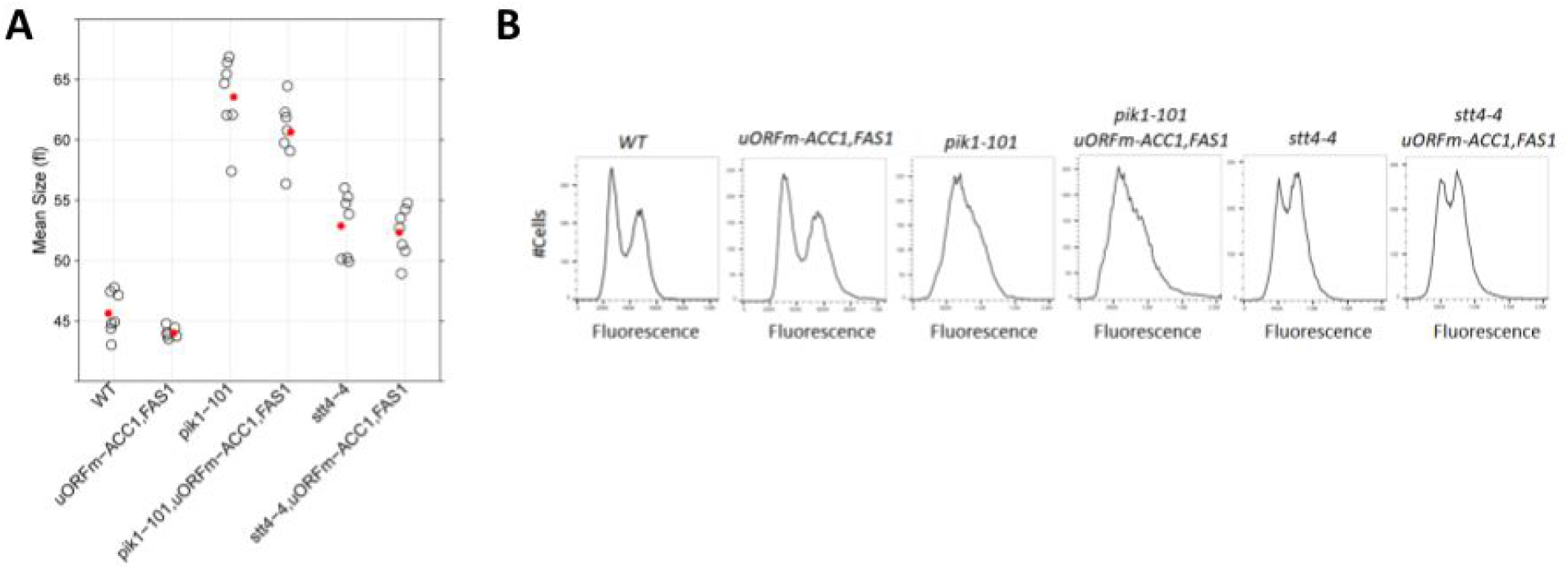
Cells lacking Pik1p or Stt4p are large (A) and have a DNA content consistent with a delay in the S phase (B). Cell size and DNA content of the indicated strains were measured at 25°C in YPD medium, as described in Materials and Methods and in Figure 4.

**FIGURE 7 - figure supplement 2.**
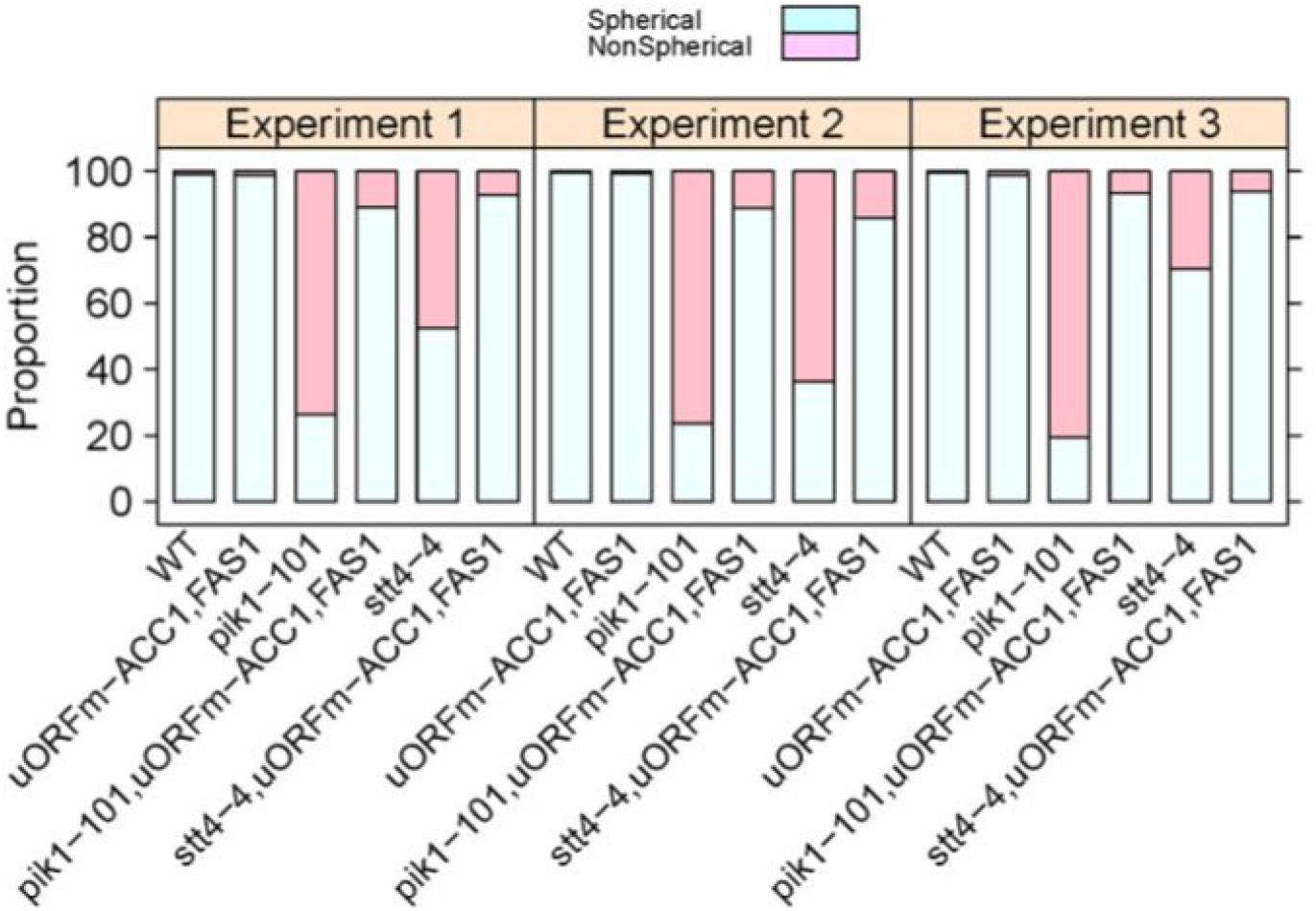
Scoring the nuclear shape of PI-4-kinase mutants, from three independent experiments.

## Notes

### Competing Interest Statement

The authors have declared no competing interest.

## REFERENCES

Al-Feel, W., S.S. Chirala, and S.J. Wakil. 1992. Cloning of the yeast FAS3 gene and primary structure of yeast acetyl-CoA carboxylase. Proc. Natl. Acad. Sci. U. S. A. 89:4534–4538. doi:10.1073/pnas.89.10.4534.

Al-Feel, W., J.C. DeMar, and S.J. Wakil. 2003. A Saccharomyces cerevisiae mutant strain defective in acetyl-CoA carboxylase arrests at the G2/M phase of the cell cycle. Proc Natl Acad Sci U A. 100:3095–100. doi:10.1073/pnas.0538069100.

Amberg, D.C., D.J. Burke, and J.N. Strathern. 2006. Yeast protein extracts. CSH Protoc. 2006:pdb. prot4152. doi:10.1101/pdb.prot4152.

Aramayo, R., and M. Polymenis. 2017. Ribosome profiling the cell cycle: lessons and challenges. Curr Genet. 63:959–964. doi:10.1007/s00294-017-0698-3.

Atilla-Gokcumen, G.E., E. Muro, J. Relat-Goberna, S. Sasse, A. Bedigian, M.L. Coughlin, S. Garcia-Manyes, and U.S. Eggert. 2014. Dividing cells regulate their lipid composition and localization. Cell. 156:428–39. doi:10.1016/j.cell.2013.12.015.

Audhya, A., M. Foti, and S.D. Emr. 2000. Distinct Roles for the Yeast Phosphatidylinositol 4-Kinases, Stt4p and Pik1p, in Secretion, Cell Growth, and Organelle Membrane Dynamics. Mol. Biol. Cell. 11:2673–2689. doi:10.1091/mbc.11.8.2673.

Baenke, F., B. Peck, H. Miess, and A. Schulze. 2013. Hooked on fat: the role of lipid synthesis in cancer metabolism and tumour development. Model Mech. 6:1353–63. doi:10.1242/dmm.011338.

Bankaitis, V.A., J.R. Aitken, A.E. Cleves, and W. Dowhan. 1990. An essential role for a phospholipid transfer protein in yeast Golgi function. Nature. 347:561–562. doi:10.1038/347561a0.

Bankaitis, V.A., D.E. Malehorn, S.D. Emr, and R. Greene. 1989. The Saccharomyces cerevisiae SEC14 gene encodes a cytosolic factor that is required for transport of secretory proteins from the yeast Golgi complex. J. Cell Biol. 108:1271–1281. doi:10.1083/jcb.108.4.1271.

Bankaitis, V.A., C.J. Mousley, and G. Schaaf. 2010. The Sec14 superfamily and mechanisms for crosstalk between lipid metabolism and lipid signaling. Trends Biochem Sci. 35:150–60. doi:10.1016/j.tibs.2009.10.008.

Barbosa, A.D., K. Lim, M. Mari, J.R. Edgar, L. Gal, P. Sterk, B.J. Jenkins, A. Koulman, D.B. Savage, M. Schuldiner, F. Reggiori, P.A. Wigge, and S. Siniossoglou. 2019. Compartmentalized Synthesis of Triacylglycerol at the Inner Nuclear Membrane Regulates Nuclear Organization. Dev. Cell. 50:755–766.e6. doi:10.1016/j.devcel.2019.07.009.

Beloribi-Djefaflia, S., S. Vasseur, and F. Guillaumond. 2016. Lipid metabolic reprogramming in cancer cells. Oncogenesis. 5:e189. doi:10.1038/oncsis.2015.49.

Blank, H.M., N. Maitra, and M. Polymenis. 2017a. Lipid biosynthesis: When the cell cycle meets protein synthesis? Cell Cycle. 16:905–906. doi:10.1080/15384101.2017.1312851.

Blank, H.M., O. Papoulas, N. Maitra, R. Garge, B.K. Kennedy, B. Schilling, E.M. Marcotte, and M. Polymenis. 2020. Abundances of transcripts, proteins, and metabolites in the cell cycle of budding yeast reveal coordinate control of lipid metabolism. Mol. Biol. Cell. 31:1069–1084. doi:10.1091/mbc.E19-12-0708.

Blank, H.M., R. Perez, C. He, N. Maitra, R. Metz, J. Hill, Y. Lin, C.D. Johnson, V.A. Bankaitis, B.K. Kennedy, R. Aramayo, and M. Polymenis. 2017b. Translational control of lipogenic enzymes in the cell cycle of synchronous, growing yeast cells. EMBO J. 36:487–502. doi:10.15252/embj.201695050.

Cameroni, E., C. De Virgilio, and O. Deloche. 2006. Phosphatidylinositol 4-phosphate is required for translation initiation in Saccharomyces cerevisiae. J. Biol. Chem. 281:38139–38149. doi:10.1074/jbc.M601060200.

Chen, M., T. Wen, H.T. Horn, V.K. Chandrahas, N. Thapa, S. Choi, V.L. Cryns, and R.A. Anderson. 2020. The nuclear phosphoinositide response to stress. Cell Cycle Georget. Tex. 19:268–289. doi:10.1080/15384101.2019.1711316.

Cleves, A.E., T.P. McGee, E.A. Whitters, K.M. Champion, J.R. Aitken, W. Dowhan, M. Goebl, and V.A. Bankaitis. 1991. Mutations in the CDP-choline pathway for phospholipid biosynthesis bypass the requirement for an essential phospholipid transfer protein. Cell. 64:789–800.

Cleves, A.E., P.J. Novick, and V.A. Bankaitis. 1989. Mutations in the SAC1 gene suppress defects in yeast Golgi and yeast actin function. J. Cell Biol. 109:2939–2950. doi:10.1083/jcb.109.6.2939.

Cocco, L., R.S. Gilmour, A. Ognibene, A.J. Letcher, F.A. Manzoli, and R.F. Irvine. 1987. Synthesis of polyphosphoinositides in nuclei of Friend cells. Evidence for polyphosphoinositide metabolism inside the nucleus which changes with cell differentiation. Biochem. J. 248:765–770. doi:10.1042/bj2480765.

Cocco, L., A.M. Martelli, R.S. Gilmour, A. Ognibene, F.A. Manzoli, and R.F. Irvine. 1989. Changes in nuclear inositol phospholipids induced in intact cells by insulin-like growth factor I. Biochem. Biophys. Res. Commun. 159:720–725. doi:10.1016/0006-291x(89)90054-5.

Collart, M.A., and S. Oliviero. 2001. Preparation of yeast RNA. Curr Protoc Mol Biol. Chapter 13:Unit 13.12. doi:10.1002/0471142727.mb1312s23.

Creanor, J., and J. Mitchison. 1979. Reduction of perturbations in leucine incorporation in synchronous cultures of Schizosaccharomyces pombe made by elutriation. J. Gen. Microbiol. 112:385–388.

Dacquay, L., A. Flint, J. Butcher, D. Salem, M. Kennedy, M. Kaern, A. Stintzi, and K. Baetz. 2017. NuA4 Lysine Acetyltransferase Complex Contributes to Phospholipid Homeostasis in Saccharomyces cerevisiae. G3 Bethesda. 7:1799–1809. doi:10.1534/g3.117.041053.

Elliott, S.G., J.R. Warner, and C.S. McLaughlin. 1979. Synthesis of ribosomal proteins during the cell cycle of the yeast Saccharomyces cerevisiae. J Bacteriol. 137:1048–50.

Fang, M., B.G. Kearns, A. Gedvilaite, S. Kagiwada, M. Kearns, M.K. Fung, and V.A. Bankaitis. 1996. Kes1p shares homology with human oxysterol binding protein and participates in a novel regulatory pathway for yeast Golgi-derived transport vesicle biogenesis. Embo J. 15:6447–59.

Fischer, M., M. Joppe, B. Mulinacci, R. Vollrath, K. Konstantinidis, P. Kötter, L. Ciccarelli, J. Vonck, D. Oesterhelt, and M. Grininger. 2020. Analysis of the co-translational assembly of the fungal fatty acid synthase (FAS). Sci. Rep. 10:895. doi:10.1038/s41598-020-57418-8.

Gao, S., K.P. Kinzig, S. Aja, K.A. Scott, W. Keung, S. Kelly, K. Strynadka, S. Chohnan, W.W. Smith, K.L.K. Tamashiro, E.E. Ladenheim, G.V. Ronnett, Y. Tu, M.J. Birnbaum, G.D. Lopaschuk, and T.H. Moran. 2007. Leptin activates hypothalamic acetyl-CoA carboxylase to inhibit food intake. Proc. Natl. Acad. Sci. U. S. A. 104:17358–17363. doi:10.1073/pnas.0708385104.

Garcia-Bustos, J.F., F. Marini, I. Stevenson, C. Frei, and M.N. Hall. 1994. PIK1, an essential phosphatidylinositol 4-kinase associated with the yeast nucleus. EMBO J. 13:2352–2361.

Giaever, G., A.M. Chu, L. Ni, C. Connelly, L. Riles, S. Veronneau, S. Dow, A. Lucau-Danila, K. Anderson, B. Andre, A.P. Arkin, A. Astromoff, M. El-Bakkoury, R. Bangham, R. Benito, S. Brachat, S. Campanaro, M. Curtiss, K. Davis, A. Deutschbauer, K.D. Entian, P. Flaherty, F. Foury, D.J. Garfinkel, M. Gerstein, D. Gotte, U. Guldener, J.H. Hegemann, S. Hempel, Z. Herman, D.F. Jaramillo, D.E. Kelly, S.L. Kelly, P. Kotter, D. LaBonte, D.C. Lamb, N. Lan, H. Liang, H. Liao, L. Liu, C. Luo, M. Lussier, R. Mao, P. Menard, S.L. Ooi, J.L. Revuelta, C.J. Roberts, M. Rose, P. Ross-Macdonald, B. Scherens, G. Schimmack, B. Shafer, D.D. Shoemaker, S. Sookhai-Mahadeo, R.K. Storms, J.N. Strathern, G. Valle, M. Voet, G. Volckaert, C.Y. Wang, T.R. Ward, J. Wilhelmy, E.A. Winzeler, Y. Yang, G. Yen, E. Youngman, K. Yu, H. Bussey, J.D. Boeke, M. Snyder, P. Philippsen, R.W. Davis, and M. Johnston. 2002. Functional profiling of the Saccharomyces cerevisiae genome. Nature. 418:387–91. doi:10.1038/nature00935.

Grabon, A., V.A. Bankaitis, and M.I. McDermott. 2019. The interface between phosphatidylinositol transfer protein function and phosphoinositide signaling in higher eukaryotes. J. Lipid Res. 60:242–268. doi:10.1194/jlr.R089730.

Hamann, B.L., and R.D. Blind. 2018. Nuclear phosphoinositide regulation of chromatin. J. Cell. Physiol. 233:107–123. doi:10.1002/jcp.25886.

Hoose, S.A., J.A. Rawlings, M.M. Kelly, M.C. Leitch, Q.O. Ababneh, J.P. Robles, D. Taylor, E.M. Hoover, B. Hailu, K.A. McEnery, S.S. Downing, D. Kaushal, Y. Chen, A. Rife, K.A. Brahmbhatt, R. Smith 3rd, and M. Polymenis. 2012. A systematic analysis of cell cycle regulators in yeast reveals that most factors act independently of cell size to control initiation of division. PLoS Genet. 8:e1002590. doi:10.1371/journal.pgen.1002590.

Huang, J., C.J. Mousley, L. Dacquay, N. Maitra, G. Drin, C. He, N.D. Ridgway, A. Tripathi, M. Kennedy, B.K. Kennedy, W. Liu, K. Baetz, M. Polymenis, and V.A. Bankaitis. 2018. A Lipid Transfer Protein Signaling Axis Exerts Dual Control of Cell-Cycle and Membrane Trafficking Systems. Dev Cell. 44:378–391.e5. doi:10.1016/j.devcel.2017.12.026.

Hunkeler, M., E. Stuttfeld, A. Hagmann, S. Imseng, and T. Maier. 2016. The dynamic organization of fungal acetyl-CoA carboxylase. Nat. Commun. 7:11196. doi:10.1038/ncomms11196.

Hurt, E.C. 1988. A novel nucleoskeletal-like protein located at the nuclear periphery is required for the life cycle of Saccharomyces cerevisiae. EMBO J. 7:4323–4334.

Ingolia, N.T., S. Ghaemmaghami, J.R. Newman, and J.S. Weissman. 2009. Genome-wide analysis in vivo of translation with nucleotide resolution using ribosome profiling. Science. 324:218–23. doi:10.1126/science.1168978.

Jenni, S., M. Leibundgut, D. Boehringer, C. Frick, B. Mikolásek, and N. Ban. 2007. Structure of fungal fatty acid synthase and implications for iterative substrate shuttling. Science. 316:254–261. doi:10.1126/science.1138248.

Johnston, G.C., J.R. Pringle, and L.H. Hartwell. 1977. Coordination of growth with cell division in the yeast Saccharomyces cerevisiae. Exp Cell Res. 105:79–98.

Kaiser, C., S. Michaelis, A. Mitchell, and Cold Spring Harbor Laboratory. 1994. Methods in yeast genetics: a Cold Spring Harbor Laboratory course manual. 1994th ed. Cold Spring Harbor Laboratory Press, Cold Spring Harbor, NY. vii, 234 p. pp.

Kim, K.H. 1997. Regulation of mammalian acetyl-coenzyme A carboxylase. Annu. Rev. Nutr. 17: 77–99. doi:10.1146/annurev.nutr.17.1.77.

Kuhajda, F.P., K. Jenner, F.D. Wood, R.A. Hennigar, L.B. Jacobs, J.D. Dick, and G.R. Pasternack. 1994. Fatty acid synthesis: a potential selective target for antineoplastic therapy. Proc Natl Acad Sci U A. 91:6379–83.

Kume, K., H. Cantwell, F.R. Neumann, A.W. Jones, A.P. Snijders, and P. Nurse. 2017. A systematic genomic screen implicates nucleocytoplasmic transport and membrane growth in nuclear size control. PLoS Genet. 13:e1006767. doi:10.1371/journal.pgen.1006767.

Lee, A.Y., R.P. St Onge, M.J. Proctor, I.M. Wallace, A.H. Nile, P.A. Spagnuolo, Y. Jitkova, M. Gronda, Y. Wu, M.K. Kim, K. Cheung-Ong, N.P. Torres, E.D. Spear, M.K.L. Han, U. Schlecht, S. Suresh, G. Duby, L.E. Heisler, A. Surendra, E. Fung, M.L. Urbanus, M. Gebbia, E. Lissina, M. Miranda, J.H. Chiang, A.M. Aparicio, M. Zeghouf, R.W. Davis, J. Cherfils, M. Boutry, C.A. Kaiser, C.L. Cummins, W.S. Trimble, G.W. Brown, A.D. Schimmer, V.A. Bankaitis, C. Nislow, G.D. Bader, and G. Giaever. 2014. Mapping the cellular response to small molecules using chemogenomic fitness signatures. Science. 344:208–211. doi:10.1126/science.1250217.

Leibundgut, M., T. Maier, S. Jenni, and N. Ban. 2008. The multienzyme architecture of eukaryotic fatty acid synthases. Curr. Opin. Struct. Biol. 18:714–725. doi:10.1016/j.sbi.2008.09.008.

Li, W., R.S. Laishram, and R.A. Anderson. 2013. The novel poly(A) polymerase Star-PAP is a signal-regulated switch at the 3’-end of mRNAs. Adv. Biol. Regul. 53:64–76. doi:10.1016/j.jbior.2012.10.004.

Li, X., M.P. Rivas, M. Fang, J. Marchena, B. Mehrotra, A. Chaudhary, L. Feng, G.D. Prestwich, and V.A. Bankaitis. 2002. Analysis of oxysterol binding protein homologue Kes1p function in regulation of Sec14p-dependent protein transport from the yeast Golgi complex. J Cell Biol. 157:63–77. doi:10.1083/jcb.200201037.

Lomakin, I.B., Y. Xiong, and T.A. Steitz. 2007. The Crystal Structure of Yeast Fatty Acid Synthase, a Cellular Machine with Eight Active Sites Working Together. Cell. 129:319–332. doi:10.1016/j.cell.2007.03.013.

Longtine, M.S., A. McKenzie 3rd, D.J. Demarini, N.G. Shah, A. Wach, A. Brachat, P. Philippsen, and J.R. Pringle. 1998. Additional modules for versatile and economical PCR-based gene deletion and modification in Saccharomyces cerevisiae. Yeast. 14:953–61. doi:10.1002/(SICI)1097-0061(199807)14:10<V;953::AID-YEA293>3.0.CO;2-U.

Mair, P., and R. Wilcox. 2016. Robust statistical methods in r using the wrs2 package. Harv. Univ.

Maitra, N., C. He, H.M. Blank, M. Tsuchiya, B. Schilling, M. Kaeberlein, R. Aramayo, B.K. Kennedy, and M. Polymenis. 2020. Translational control of one-carbon metabolism underpins ribosomal protein phenotypes in cell division and longevity. eLife. 9:e53127. doi:10.7554/eLife.53127.

Meseroll, R.A., and O. Cohen-Fix. 2016. The Malleable Nature of the Budding Yeast Nuclear Envelope: Flares, Fusion, and Fenestrations. J. Cell. Physiol. 231:2353–2360. doi:10.1002/jcp.25355.

Mousley, C.J., P. Yuan, N.A. Gaur, K.D. Trettin, A.H. Nile, S.J. Deminoff, B.J. Dewar, M. Wolpert, J.M. Macdonald, P.K. Herman, A.G. Hinnebusch, and V.A. Bankaitis. 2012. A sterol-binding protein integrates endosomal lipid metabolism with TOR signaling and nitrogen sensing. Cell. 148:702–15. doi:10.1016/j.cell.2011.12.026.

Nile, A.H., A. Tripathi, P. Yuan, C.J. Mousley, S. Suresh, I.M. Wallace, S.D. Shah, D.T. Pohlhaus, B. Temple, C. Nislow, G. Giaever, A. Tropsha, R.W. Davis, R.P. St Onge, and V.A. Bankaitis. 2014. PITPs as targets for selectively interfering with phosphoinositide signaling in cells. Nat Chem Biol. 10:76–84. doi:10.1038/nchembio.1389.

Novick, P., C. Field, and R. Schekman. 1980. Identification of 23 complementation groups required for post-translational events in the yeast secretory pathway. Cell. 21:205–215. doi:10.1016/0092-8674(80)90128-2.

Omura, S. 1976. The antibiotic cerulenin, a novel tool for biochemistry as an inhibitor of fatty acid synthesis. Bacteriol Rev. 40:681–97.

Price, A.C., K.-H. Choi, R.J. Heath, Z. Li, S.W. White, and C.O. Rock. 2001. Inhibition of β-Ketoacyl-Acyl Carrier Protein Synthases by Thiolactomycin and Cerulenin STRUCTURE AND MECHANISM. J. Biol. Chem. 276:6551–6559. doi:10.1074/jbc.M007101200.

Pringle, J.R., Hartwell, L.H. 1981. The Saccharomyces cerevisiae Cell Cycle. In The Molecular and Cellular Biology of the Yeast Saccharomyces. Cold Spring Harbor Laboratory Press. 97–142.

Ricciardi, S., N. Manfrini, R. Alfieri, P. Calamita, M.C. Crosti, S. Gallo, R. Müller, M. Pagani, S. Abrignani, and S. Biffo. 2018. The Translational Machinery of Human CD4+ T Cells Is Poised for Activation and Controls the Switch from Quiescence to Metabolic Remodeling. Cell Metab. 28:895–906.e5. doi:10.1016/j.cmet.2018.08.009.

Rivas, M.P., B.G. Kearns, Z. Xie, S. Guo, M.C. Sekar, K. Hosaka, S. Kagiwada, J.D. York, and V.A. Bankaitis. 1999. Pleiotropic alterations in lipid metabolism in yeast sac1 mutants: relationship to “bypass Sec14p” and inositol auxotrophy. Mol Biol Cell. 10:2235–50.

Santos-Rosa, H., J. Leung, N. Grimsey, S. Peak-Chew, and S. Siniossoglou. 2005. The yeast lipin Smp2 couples phospholipid biosynthesis to nuclear membrane growth. Embo J. 24:1931–41. doi:10.1038/sj.emboj.7600672.

Scaglia, N., S. Tyekucheva, G. Zadra, C. Photopoulos, and M. Loda. 2014. De novo fatty acid synthesis at the mitotic exit is required to complete cellular division. Cell Cycle. 13:859–68. doi:10.4161/cc.27767.

Schaaf, G., E.A. Ortlund, K.R. Tyeryar, C.J. Mousley, K.E. Ile, T.A. Garrett, J. Ren, M.J. Woolls, C.R. Raetz, M.R. Redinbo, and V.A. Bankaitis. 2008. Functional anatomy of phospholipid binding and regulation of phosphoinositide homeostasis by proteins of the sec14 superfamily. Mol Cell. 29:191–206. doi:10.1016/j.molcel.2007.11.026.

Schneiter, R., M. Hitomi, A.S. Ivessa, E.V. Fasch, S.D. Kohlwein, and A.M. Tartakoff. 1996. A yeast acetyl coenzyme A carboxylase mutant links very-long-chain fatty acid synthesis to the structure and function of the nuclear membrane-pore complex. Mol Cell Biol. 16:7161–72.

Shiber, A., K. Döring, U. Friedrich, K. Klann, D. Merker, M. Zedan, F. Tippmann, G. Kramer, and B. Bukau. 2018. Cotranslational assembly of protein complexes in eukaryotes revealed by ribosome profiling. Nature. 561:268–272. doi:10.1038/s41586-018-0462-y.

Shulman, R.W., L.H. Hartwell, and J.R. Warner. 1973. Synthesis of ribosomal proteins during the yeast cell cycle. J Mol Biol. 73:513–25.

Siniossoglou, S. 2013. Phospholipid metabolism and nuclear function: roles of the lipin family of phosphatidic acid phosphatases. Biochim Biophys Acta. 1831:575–81. doi:10.1016/j.bbalip.2012.09.014.

Soma, S., K. Yang, M.I. Morales, and M. Polymenis. 2014. Multiple metabolic requirements for size homeostasis and initiation of division in Saccharomyces cerevisiae. Microb Cell. 1:256–266. doi:10.15698/mic2014.08.160.

Storck, E.M., C. Ozbalci, and U.S. Eggert. 2018. Lipid Cell Biology: A Focus on Lipids in Cell Division. Annu Rev Biochem. 87:839–869. doi:10.1146/annurev-biochem-062917-012448.

Strahl, T., H. Hama, D.B. DeWald, and J. Thorner. 2005. Yeast phosphatidylinositol 4-kinase, Pik1, has essential roles at the Golgi and in the nucleus. J. Cell Biol. 171:967–979. doi:10.1083/jcb.200504104.

Takemoto, A., S.A. Kawashima, J.-J. Li, L. Jeffery, K. Yamatsugu, O. Elemento, and P. Nurse. 2016. Nuclear envelope expansion is crucial for proper chromosomal segregation during a closed mitosis. J. Cell Sci. 129:1250–1259. doi:10.1242/jcs.181560.

Walters, A.D., K. Amoateng, R. Wang, J.-H. Chen, G. McDermott, C.A. Larabell, O. Gadal, and O. Cohen-Fix. 2018. Nuclear envelope expansion in budding yeast is independent of cell growth and does not determine nuclear volume. Mol. Biol. Cell. 30:131–145. doi:10.1091/mbc.E18-04-0204.

Walters, A.D., A. Bommakanti, and O. Cohen-Fix. 2012. Shaping the nucleus: factors and forces. J Cell Biochem. 113:2813–21. doi:10.1002/jcb.24178.

Walters, A.D., C.K. May, E.S. Dauster, B.P. Cinquin, E.A. Smith, X. Robellet, D. D’Amours, C.A. Larabell, and O. Cohen-Fix. 2014. The yeast polo kinase Cdc5 regulates the shape of the mitotic nucleus. CurrBiol. 24:2861–7. doi:10.1016/j.cub.2014.10.029.

Wang, Y., C.J. Mousley, M.G. Lete, and V.A. Bankaitis. 2019. An equal opportunity collaboration between lipid metabolism and proteins in the control of membrane trafficking in the trans-Golgi and endosomal systems. Curr Opin Cell Biol. 59:58–72. doi:10.1016/j.ceb.2019.03.012.

Webster, M.T., J.M. McCaffery, and O. Cohen-Fix. 2010. Vesicle trafficking maintains nuclear shape in Saccharomyces cerevisiae during membrane proliferation. J. Cell Biol. 191:1079–1088. doi:10.1083/jcb.201006083.

Wenz, P. 2001. A downstream regulatory element located within the coding sequence mediates autoregulated expression of the yeast fatty acid synthase gene FAS2 by the FAS1 gene product. Nucleic Acids Res. 29:4625–4632. doi:10.1093/nar/29.22.4625.

Wilcox, R.R. 2011. Introduction to robust estimation and hypothesis testing. Academic press.

Witkin, K.L., Y. Chong, S. Shao, M.T. Webster, S. Lahiri, A.D. Walters, B. Lee, J.L. Koh, W.A. Prinz, B.J. Andrews, and O. Cohen-Fix. 2012. The budding yeast nuclear envelope adjacent to the nucleolus serves as a membrane sink during mitotic delay. Curr Biol. 22:1128–33. doi:10.1016/j.cub.2012.04.022.

Xie, Z., M. Fang, and V.A. Bankaitis. 2001. Evidence for an intrinsic toxicity of phosphatidylcholine to Sec14p-dependent protein transport from the yeast Golgi complex. Mol. Biol. Cell. 12:1117–1129. doi:10.1091/mbc.12.4.1117.

Xu, Z., W. Wei, J. Gagneur, F. Perocchi, S. Clauder-Münster, J. Camblong, E. Guffanti, F. Stutz, W. Huber, and L.M. Steinmetz. 2009. Bidirectional promoters generate pervasive transcription in yeast. Nature. 457:1033–1037. doi:10.1038/nature07728.

York, J.D., A.R. Odom, R. Murphy, E.B. Ives, and S.R. Wente. 1999. A phospholipase C-dependent inositol polyphosphate kinase pathway required for efficient messenger RNA export. Science. 285:96–100. doi:10.1126/science.285.5424.96.

Zach, R., and M. Prevorovsky. 2018. The phenomenon of lipid metabolism “cut” mutants. Yeast. 35:631–637. doi:10.1002/yea.3358.

