## Supplementary material for "Translational control of lipogenesis links protein synthesis and phosphoinositide signaling with nuclear division": Figure 1 - source data 1

Independent experiments used in the quantification of Fas1-TAP from asynchronous cultures of wild type and *uORFm-ACC1,FAS1* cells growing in media with glucose or glycerol as the carbon source. Pgk1p is the loading control. Experiments are grouped by rows with the description at the top of each column. Blots were digitally recorded with the Amersham™ Imager 600. The raw images were processed by ImageJ using the 'subtract background' and protein levels were measured by the 'measure' tool.

Experiment 1

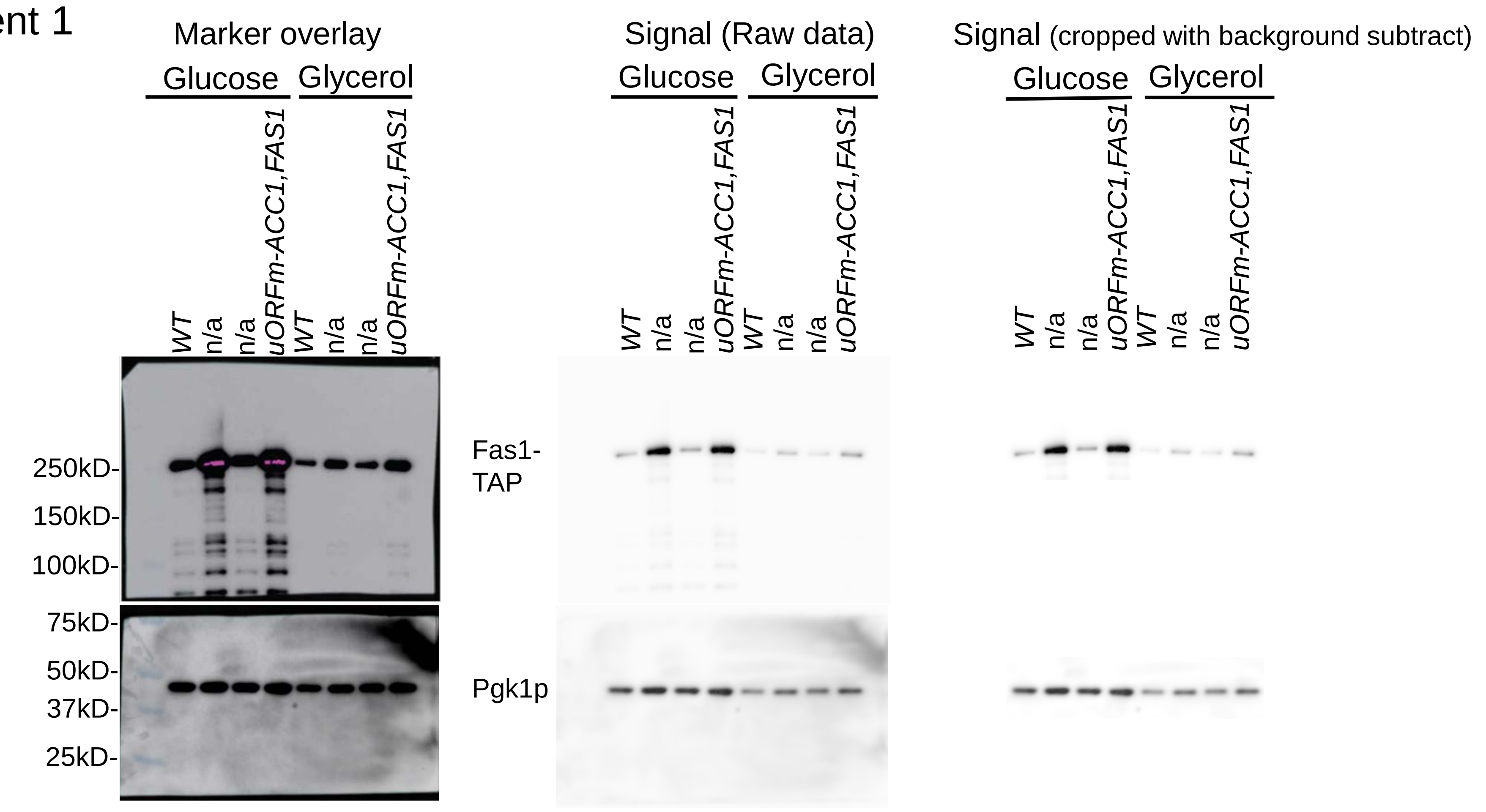


Experiment 2

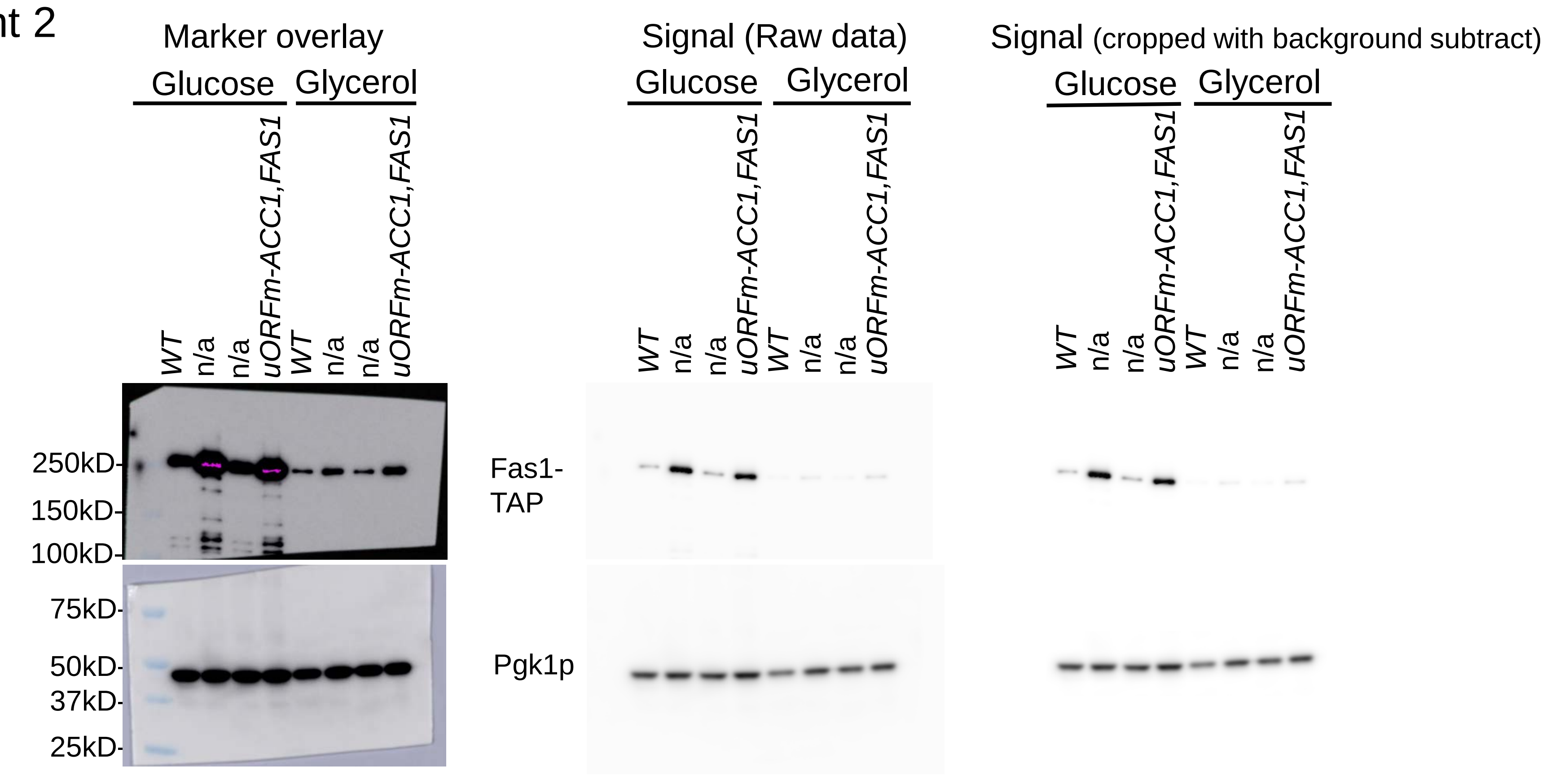


Experiment 3

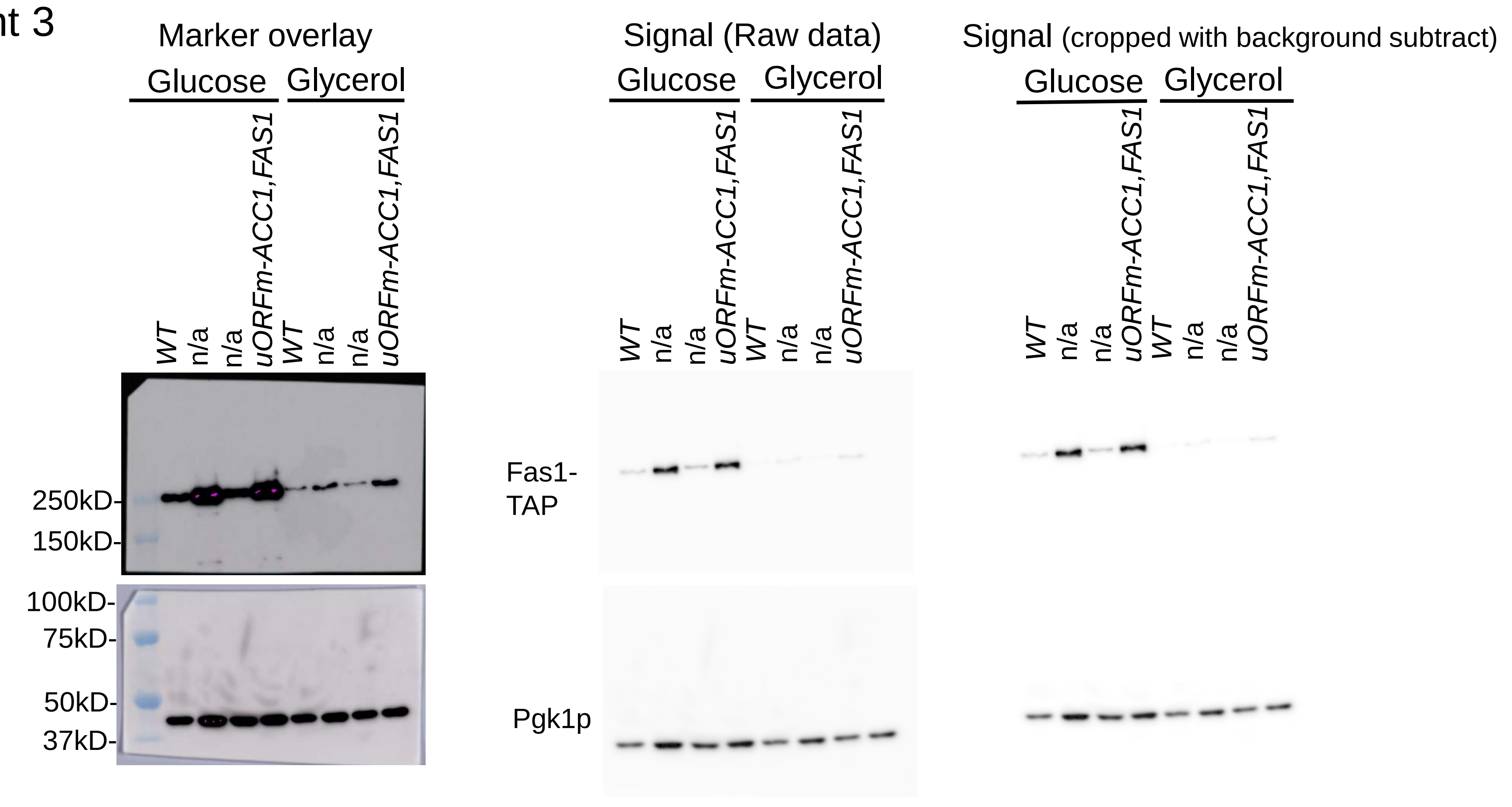


Experiment 4

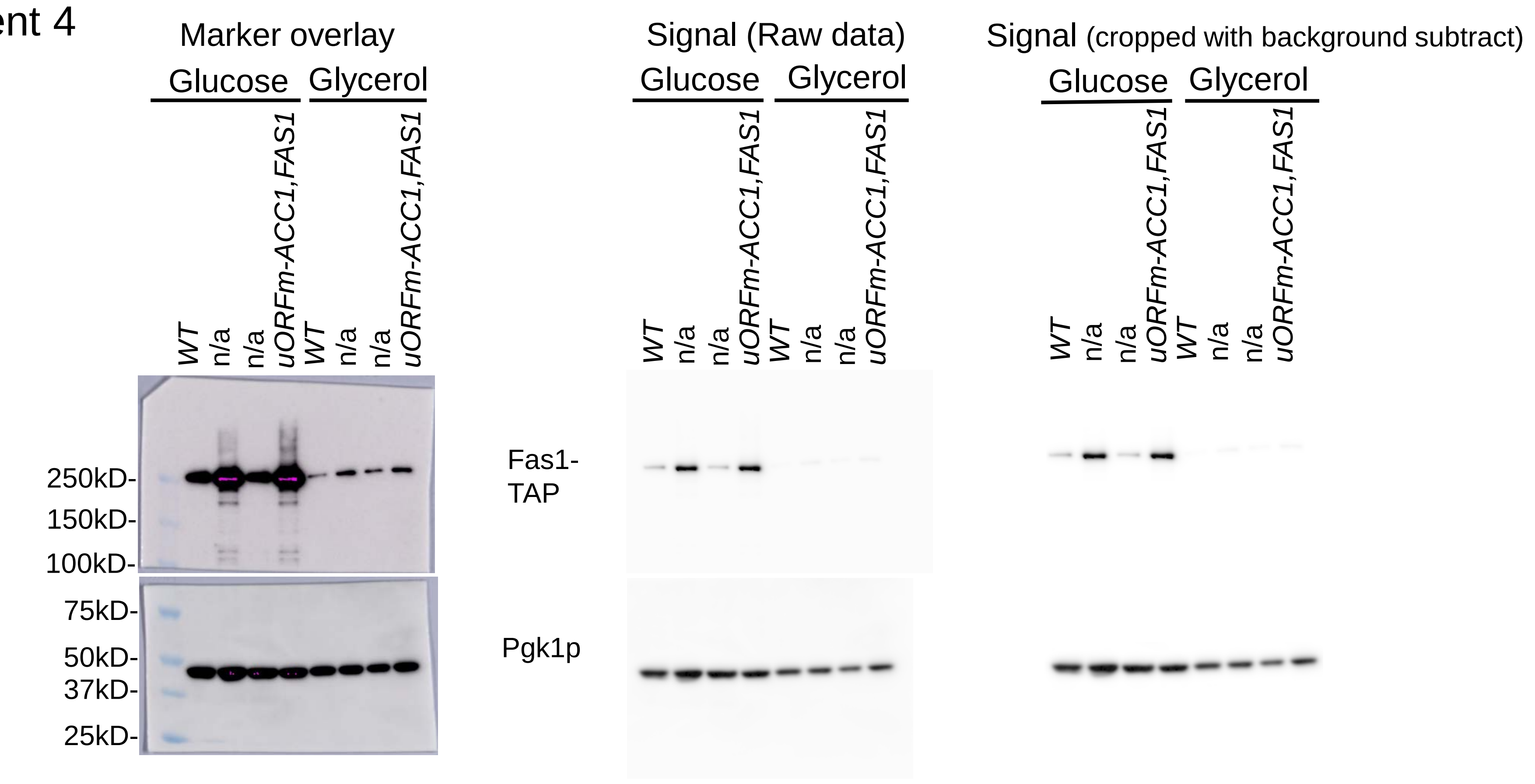


Experiment 5

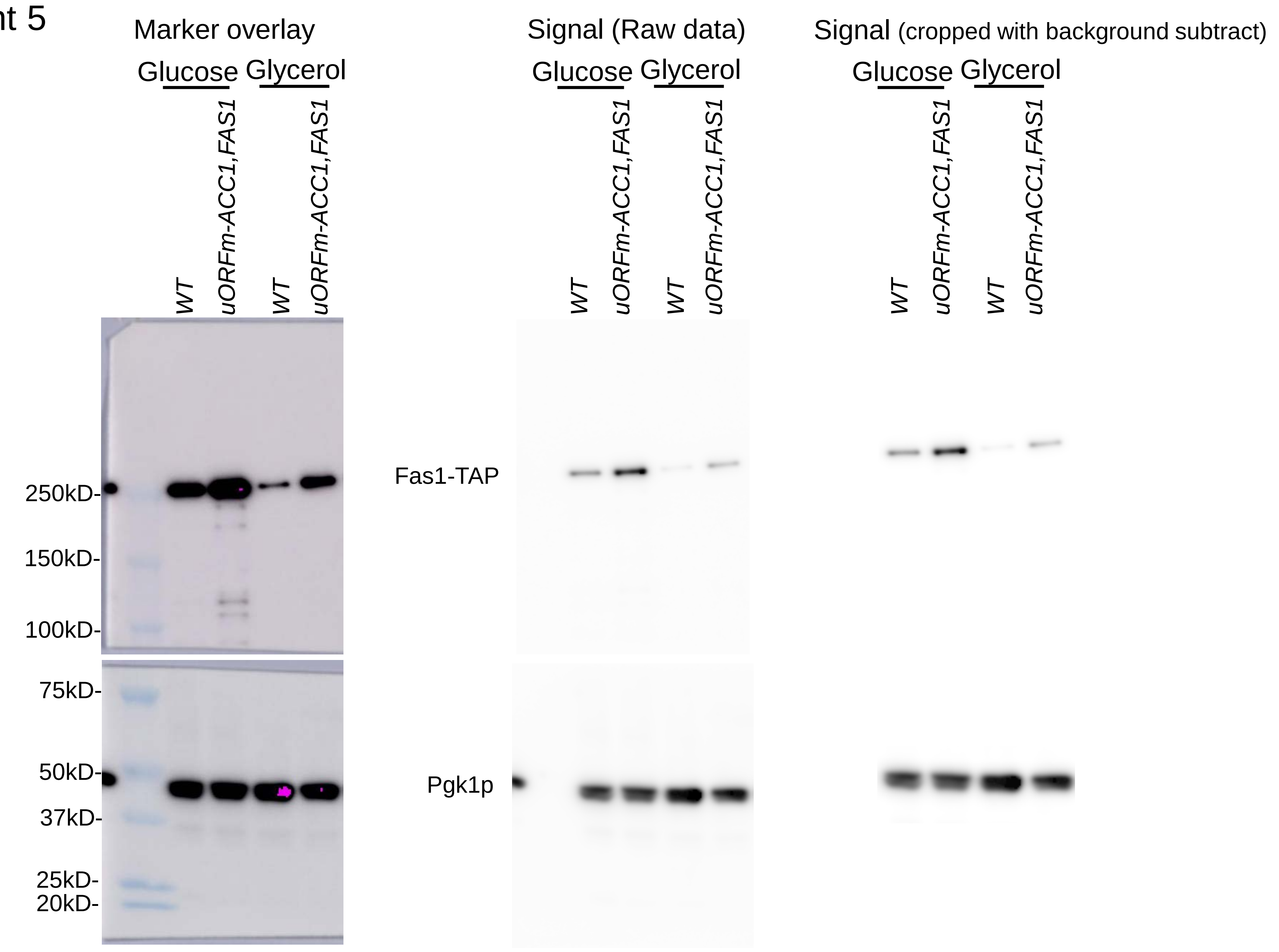
