## Supplementary material for "Translational control of lipogenesis links protein synthesis and phosphoinositide signaling with nuclear division": Figure 1 - source data 2

*FAS1* and *UBC6* from asynchronous cultures growing in media with either glucose or glycerol as carbon source were amplified and detected using FAS1-FAM probe (blue) and UBC6-VIC probe (green). In the ddPCR System, the QuantaSoft™ Software determines the numbers of droplets that are positive (blue/green colored circles in the left panel) and negative (grey circles in the left panel) for each fluorophore in each sample. The fraction of positive droplets is then fitted to a Poisson distribution to determine the absolute starting copy number in units of copies/µl input sample.

Experiment 1

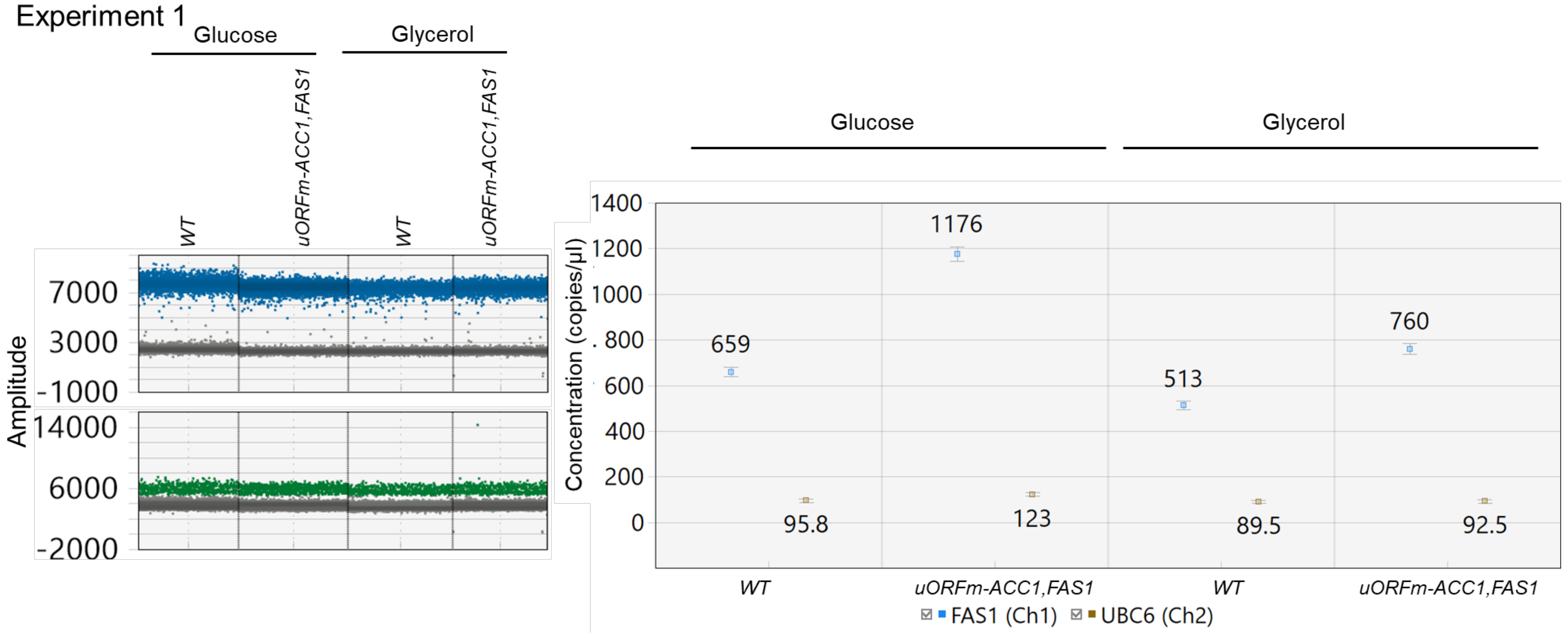


Experiment 2

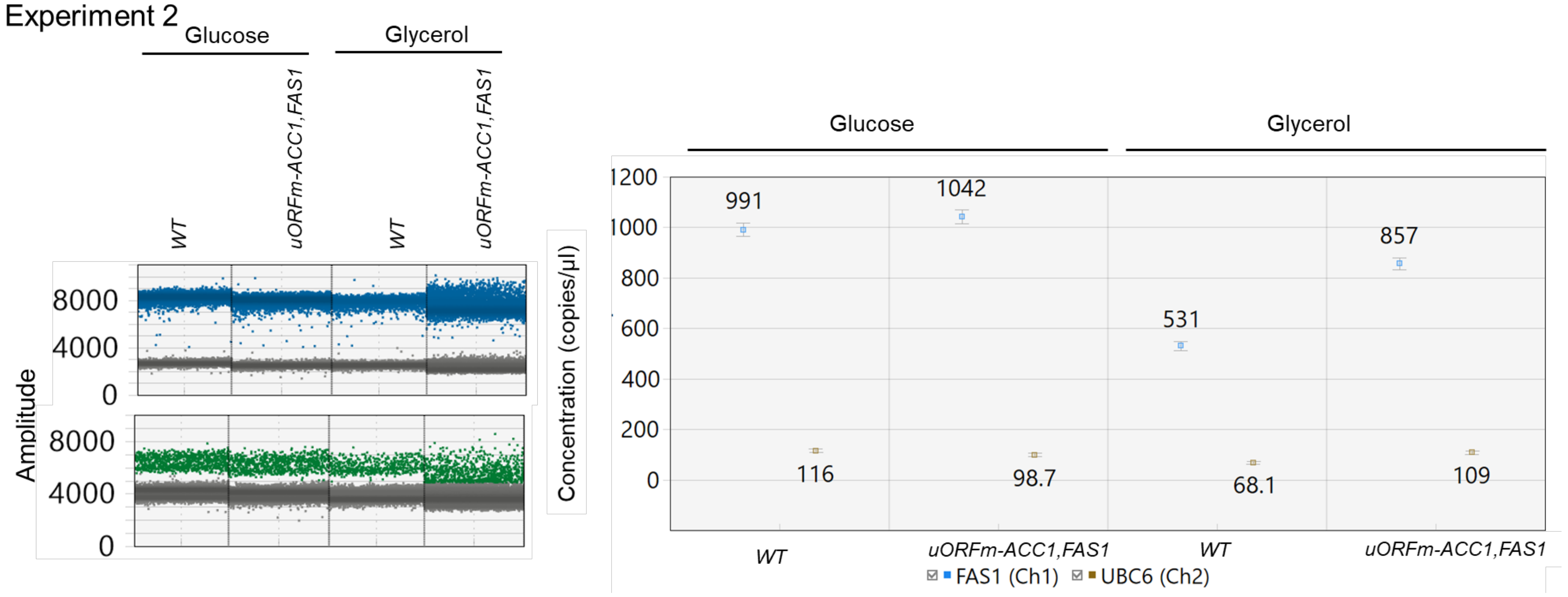


Experiment 3

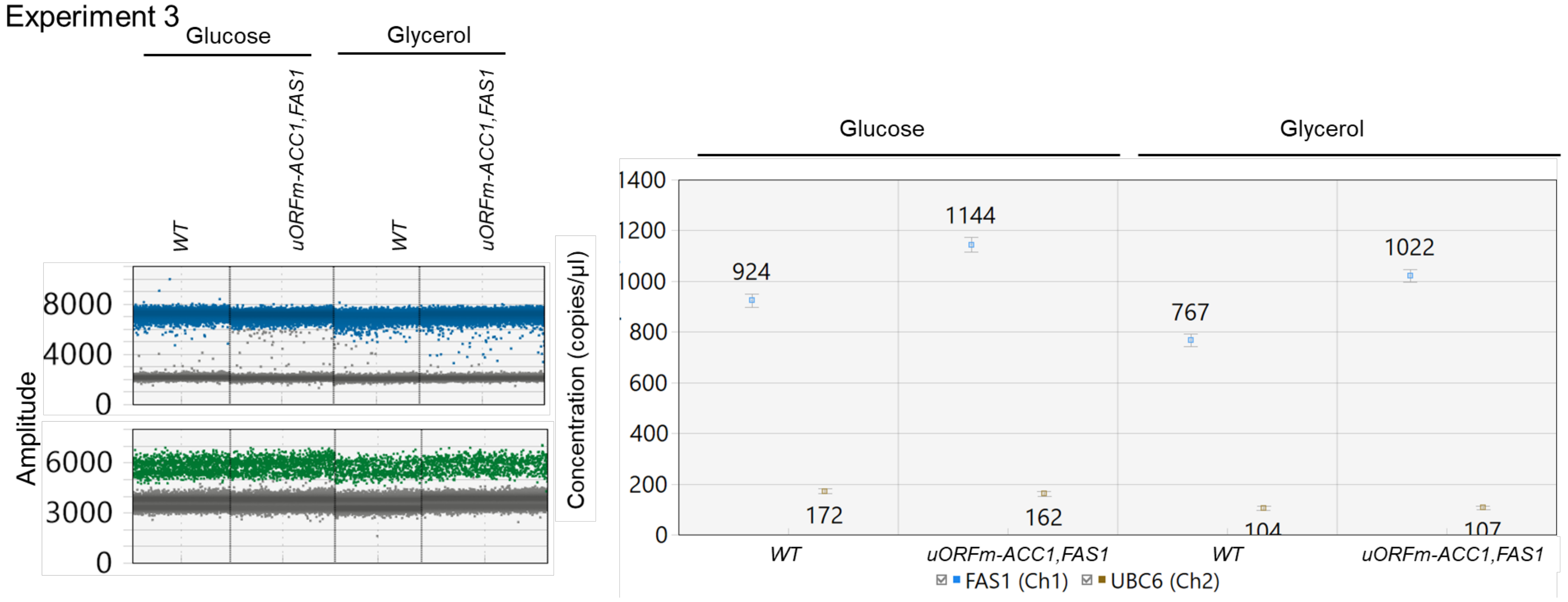


Experiment 4

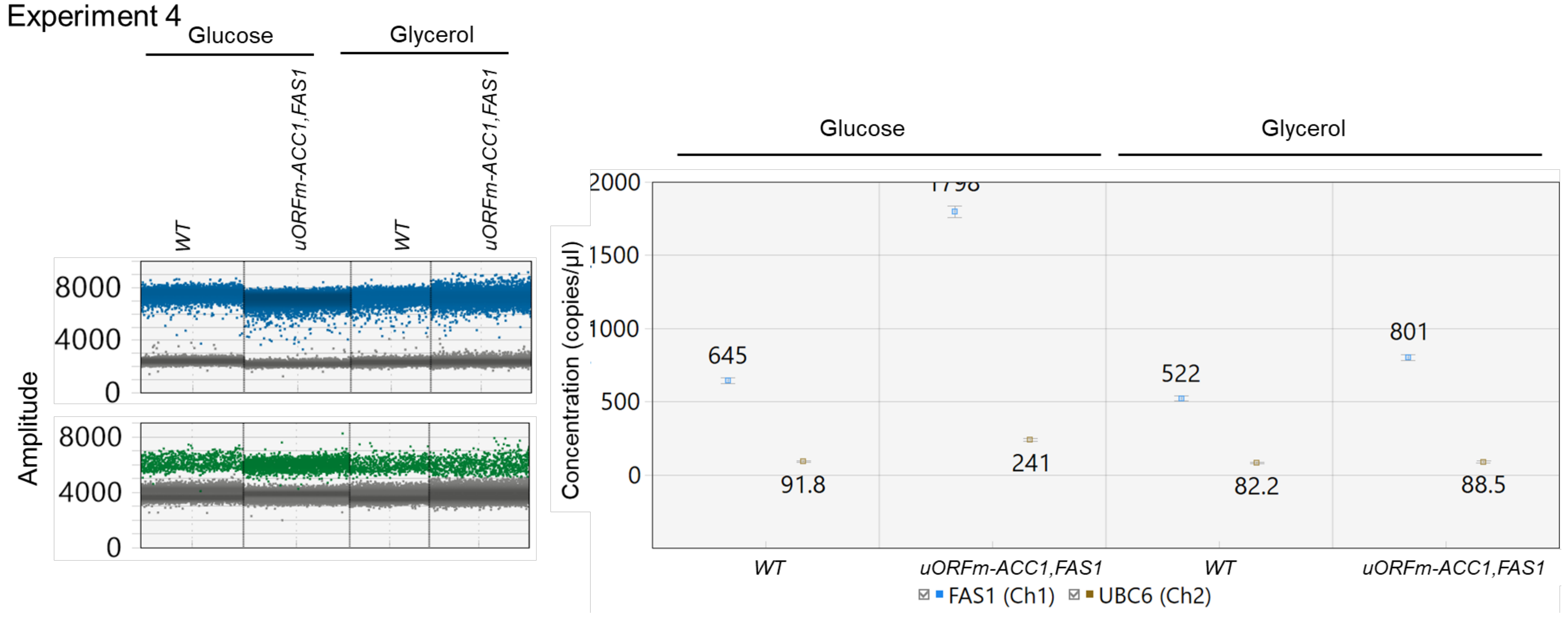
