## Supplementary material for "Translational control of lipogenesis links protein synthesis and phosphoinositide signaling with nuclear division": Figure 1 - source data 3

Independent experiments used in the quantification of Fas1-TAP levels through a synchronous cell cycle of wild type cells growing in standard YPD medium. Pgk1p is the loading control. Experiments are grouped by rows with the description at the top of each column. Blots were digitally recorded by the Amersham™ Imager 600. The raw images were processed by ImageJ using the 'subtract background' tool and protein levels were measured by the 'measure' tool.

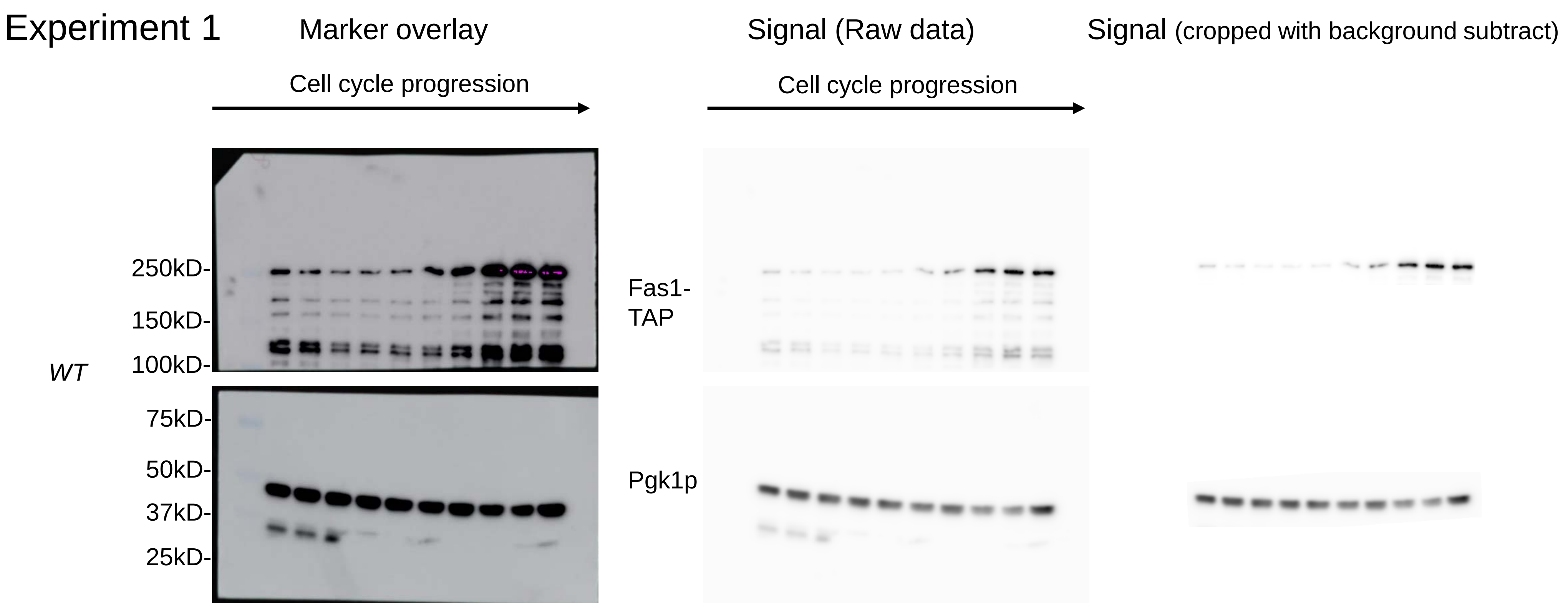


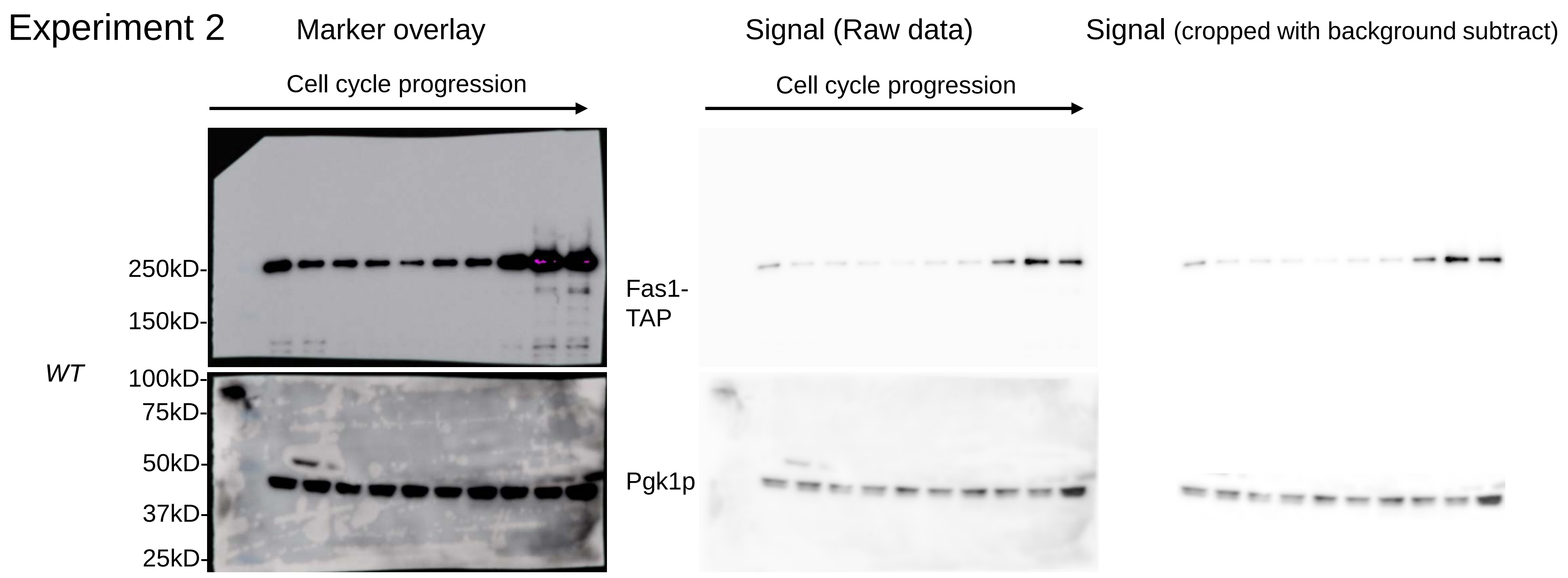


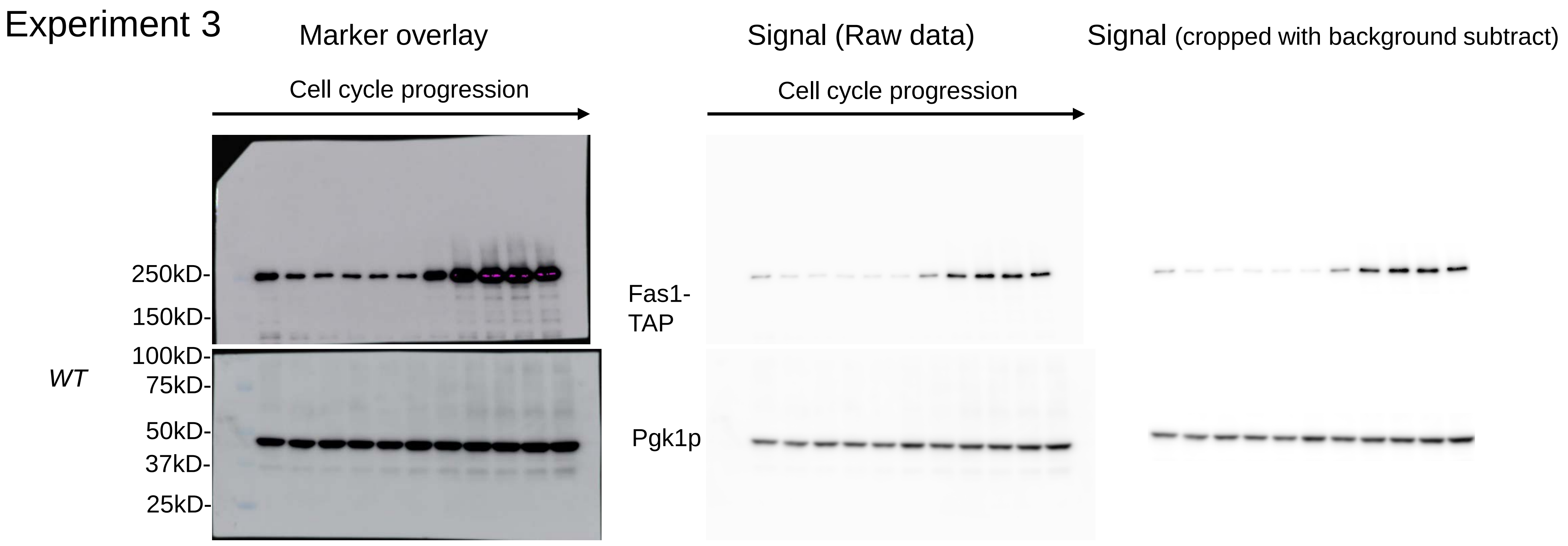


Independent experiments used in the quantification of Fas1-TAP levels through a synchronous cell cycle of *uORFm-ACC1,FAS1* cells growing in standard YPD medium. Pgk1p is the loading control. Experiments are grouped by rows with the description at the top of each column. Blots were digitally recorded by the Amersham™ Imager 600. The raw images were processed by ImageJ using the 'subtract background' tool and protein levels were measured by the 'measure' tool.

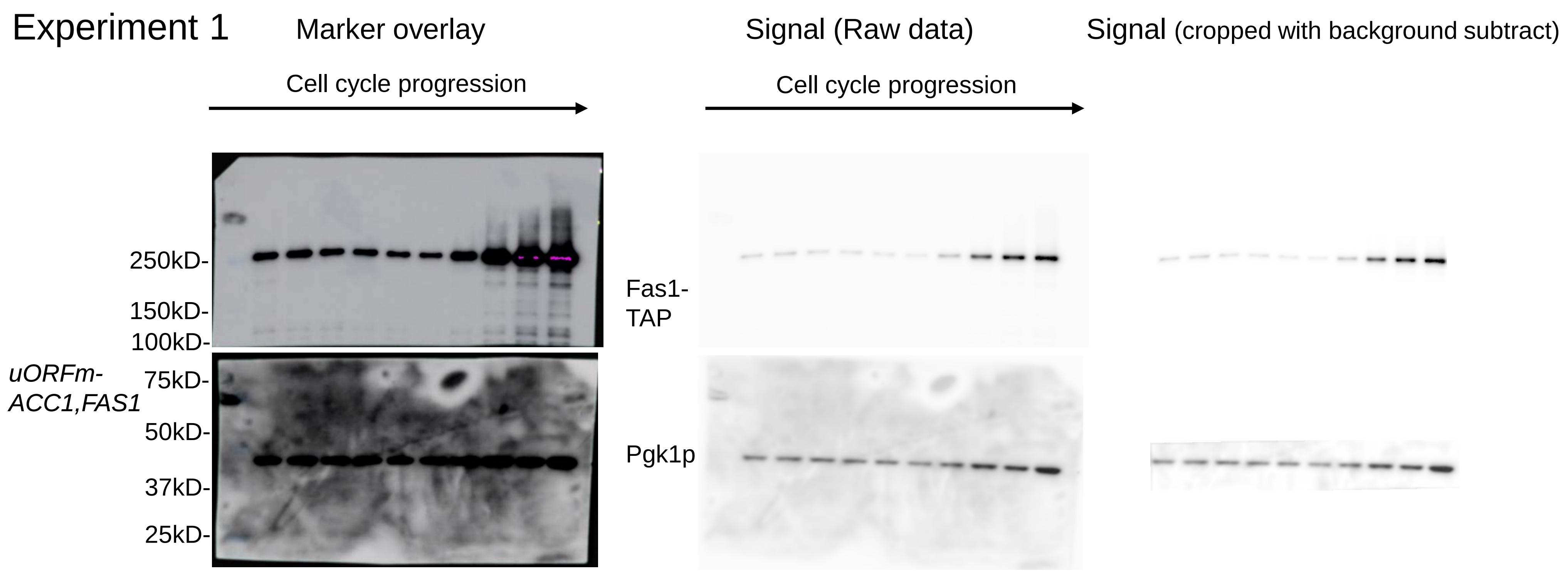


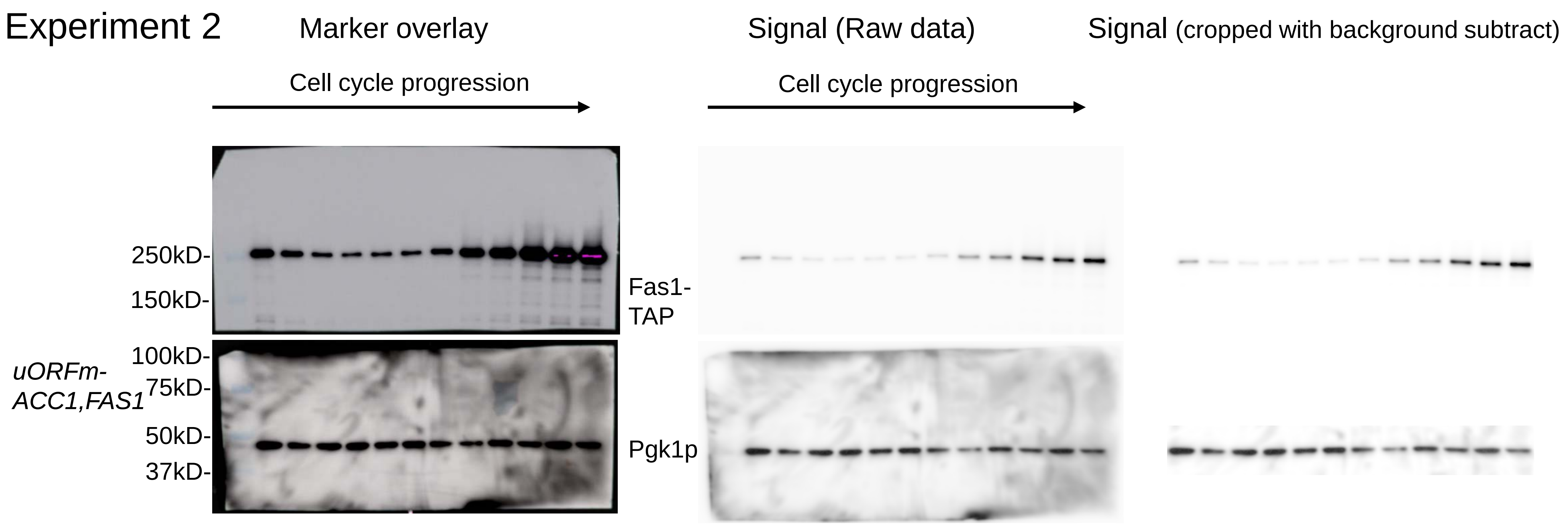


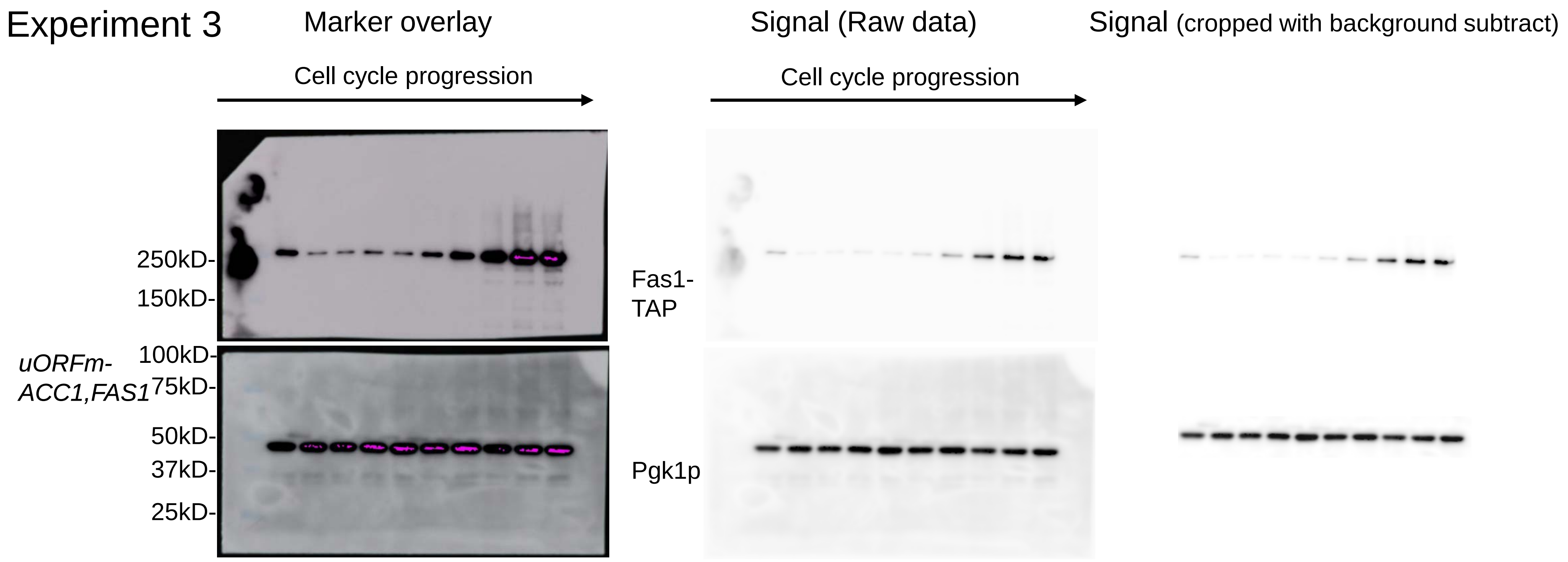
